# Delayed activation of the DNA replication licensing system in Lgr5(+) intestinal stem cells

**DOI:** 10.1101/177477

**Authors:** T.D. Carroll, I.P. Newton, Y. Chen, J.J. Blow, I. Näthke

## Abstract

During late mitosis and early G_1_, replication origins are licensed for replication by binding to double hexamers of MCM2-7. Here, we investigate how licensing and proliferative commitment are coupled in the small-intestinal epithelium. We developed a method for identifying cells in intact tissue containing DNA-bound MCM2-7. Interphase cells above the transit-amplifying compartment had no DNA-bound MCM2-7, but still expressed MCM2-7 protein, suggesting that licensing is inhibited immediately upon differentiation. Strikingly, we found most proliferative Lgr5(+) stem cells are in an unlicensed state. This suggests that the elongated cell-cycle of intestinal stem-cells is caused by an increased G_1_ length, characterised by dormant periods with unlicensed origins. Significantly, the unlicensed state is lost In *Apc* mutant epithelium, which lacks a functional restriction point, causing licensing immediately upon G_1_ entry. We propose that the unlicensed G_1_ of intestinal stem cells creates a temporal window when proliferative fate decisions can be made.

## INTRODUCTION

Cell division is necessary for adult tissue homeostasis. It allows for the replacement of aged or damaged cells and the provision of specialised cells critical for tissue function. The decision to proliferate is crucial, especially for stem cells, which produce daughter cells that either maintain a stem cell fate or differentiate to produce specialised cells. The rapidly-renewing intestinal epithelium replenishes its cellular content every 4-5 days. This high turnover rate is maintained primarily by Lgr5(+) intestinal stem cells in the crypt base, thought to be continually proliferative (Basak et al., 2014) as confirmed by proteomic and transcriptomic analysis (Munoz et al., 2012). There is also a quiescent stem cell-population that can re-engage with the cell-cycle to repopulate the Lgr5(+) cell population if it becomes depleted. These quiescent stem cells reside at the +4 position and constitute a subset of Lgr5(+) cells and are immature secretory lineage precursors (Buczacki et al., 2013). Lgr5(+) stem cells can divide to form transit-amplifying (TA) cells, which undergo several rounds of cell division before differentiating and losing proliferative competency (Potten and Loeffler, 1990).

How proliferative fate decisions are governed in stem cells and transit-amplifying cells is not understood. Lineage tracing studies suggest that in homeostatic intestinal tissue only 5-7 intestinal stem cells are ‘active’ out of the 12-16 Lgr5(+) cells present in the crypt base (Baker et al., 2014, Kozar et al., 2013). Interestingly, Lgr5(+) cells have a significantly longer cell-cycle than transit-amplifying cells (Schepers et al., 2011). The functional significance of the prolonged cell-cycle time of Lgr5(+) stem cells is currently unknown, but suggests active regulation of cell-cycle progression and proliferative fate commitment.

Proliferative fate decisions are typically visualised by detecting markers that are present in all cell-cycle phases, and only distinguish proliferative from quiescent cells. Visualising the incorporation of labelled nucleosides such as BrdU or EdU marks cells in S-phase. The limitation of these methods is that they cannot discriminate early proliferative fate decisions made during the preceding mitosis, or in the early stages of G_1_. DNA replication in S phase depends on origin licensing, involving the regulated loading of minichromosome maintenance 2-7 (MCM2-7) complexes onto origins of DNA replication (reviewed in (Blow and Hodgson, 2002, Champeris Tsaniras et al., 2014)). During S phase, DNA-bound MCM2-7 hexamers are activated to form the catalytic core of the DNA helicase as part of the CMG (Cdc45, MCM2-7, G_1_NS) complex (Moyer et al., 2006, Ilves et al., 2010, Makarova et al., 2012). Replication licensing is thought to occur from late mitosis throughout G_1_ until passage through the restriction point (Dimitrova et al., 2002, Haland et al., 2015, Namdar and Kearsey, 2006, Symeonidou et al., 2013). Correspondingly, insufficient origin licensing directly limits the ability to progress past the restriction point causing cell cycle arrest (Alver et al., 2014, Liu et al., 2009, Shreeram et al., 2002). When functional, this licensing-checkpoint can delay S-phase if an insufficient amount of origins have been licensed.

When cells enter G_0_, MCM2-7 proteins are downregulated and degraded, primarily via E2F-mediated transcriptional control of MCM2-7, Cdc6 and Cdt1 (Leone et al., 1998, Ohtani et al., 1999, Williams et al., 1998). This prevents terminally differentiated cells from re-entering the cell cycle. In mammalian cells, artificial induction of quiescence through contact inhibition leads to gradual downregulation of Cdc6 and MCM2-7 over several days (Kingsbury et al., 2005). These features have led to the suggestion that quiescence can be defined by an unlicensed state (Blow and Hodgson, 2002). Equally, the licensing status can define a different restriction point that signals proliferative fate commitment at the end of mitosis and in early G_1_, independently of the Rb/E2F restriction point.

The dynamics of replication licensing in the intricate cellular hierarchy of a complex, rapidly renewing adult tissue, is not understood. Therefore, we investigated the licensing system in the intestinal epithelium, aiming to understand dynamics of early cell-cycle commitment in stem and transit-amplifying cells and during terminal differentiation.

## MATERIALS AND METHODS

### Mice

All experiments were performed under UK home office guidelines. CL57BL/6 (Wild-type), *R26-rtTA Col1A1-H2B-GFP* (H2B-GFP), Lgr5-EGFP-IRES-creERT2 (Lgr5^GFP/+^) and Apc^Min/+^ mice were sacrificed by cervical dislocation or CO_2_ asphyxiation. Fucci2aR mice (Mort et al., 2014) were a kind Gift of Dr Richard Mort, University of Edinburgh.

### Tissue preparation: Whole small intestine

Dissected pieces of adult mouse small-intestine were washed briefly in PBS and then fixed in 4% PFA for 3 hours, 4°C. Intestines were cut into 2×2 cm^2^ pieces and fixed overnight in 4% PFA, 4°C. Tissue was embedded in 3% low melting temperature agarose and cut into 200 μm sections using a Vibratome (Leica). Sections were washed in PBS, permeabilised with 2% Triton-X100 for 2 hours and incubated with Blocking Buffer (1% BSA, 3% Normal Goat serum, 0.2% Triton-X100 in PBS) for 2 hours, 4°C. Tissue was incubated in Working Buffer (0.1% BSA, 0.3% Normal Goat Serum, 0.2% Triton-X100 in PBS) containing primary antibody, Mcm2 (Cell Signalling, 1:500), for 48 hours, 4°C. Sections were washed 5x with Working Buffer prior to 48 hour incubation with secondary antibodies diluted in Working Buffer: Alexafluor™ conjugated goat anti-rabbit (1:500, Molecular Probes) plus 5 pg/ml Hoechst 33342 and Alexafluor™ conjugated Phalloidin (1:150, Molecular Probes). Sections were mounted on coverslips in Prolong Gold between 2×120 μm spacers.

### Tissue preparation: Isolating and staining crypts

Small intestines were dissected, washed in PBS and opened longitudinally. Villi were removed by repeated (up to 10 times) scraping of the luminal surface with a coverslip. Tissue was washed in PBS, incubated in 30 mM EDTA (25 minutes, 4°C) and crypts isolated by vigorous shaking in PBS. Crypt suspensions were centrifuged (fixed rotor, 88 RCF, 4°C) and the pellet washed twice in cold PBS. Crypts were fixed in 4% PFA (30min, room temperature), permeabilized in 1% Triton-X100 (1 hour, room temperature) and blocked in Blocking Buffer (2 hours, 4°C). Crypts were incubated with primary antibodies diluted in Working Buffer: Mcm2 (Cell Signalling, 1:500), phospo-HistoneH3 (Abcam, 1:500), Ki67 (Abcam ab15580, 1:250), aGFP (Abcam, 1:500), washed 5x with Working Buffer before overnight incubation with secondary antibodies diluted in Working buffer: Alexafluor™ conjugated goat anti-mouse or anti-rabbit (1:500, Molecular Probes) or stains: Rhodamine labelled Ulex Europaeus Agglutinin I (UEA, 1:500), 5 μg/ml Hoechst 33342 or Alexafluor™ conjugated Phalloidin (1:150), at 4°C. Crypts were mounted directly on slides in Prolong Gold, overnight.

### CSK extraction of isolated crypts

Soluble proteins were extracted from crypts isolated as described above by incubation with CSK extraction buffer (10 mM HEPES, 100 mM NaCl, 3 mM MgCl_2_, 1 mM EGTA, 300 mM sucrose, 0.2% TritonX-100, 1 mM DTT, 2% BSA) supplemented with protease inhibitors (PMSF, Pepstatin, Leupeptin, Cystatin, Na_3_VO_4_, NaF, aprotinin) for 20 minutes on ice prior to fixation. Crypts were then fixed with 4% PFA and processed for imaging as described above.

### H2B-GFP label retention

H2B-GFP expression in transgenic *R26-rtTA Col1A1-H2B-GFP* mice was induced by replacing normal drinking water with 5% sucrose water supplemented with 2 mg/ml doxycycline. After 7 days, doxycycline water was replaced with normal drinking water. Subsequently, mice were sacrificed after 7 days.

### EdU incorporation and detection

Mice were injected intraperitoneally with 100 μg EdU (Invitrogen) prepared in 200 μl sterile PBS. Mice were sacrificed 1 hour or 17 hours post induction. For organoids, 10 μM EdU was included in crypt media for 1 hour before harvesting. EdU was detected by Click-it chemistry, by incubation in EdU working buffer (1.875 μM Alexafluor 488 azide (Invitrogen), 2 mM CuSO_4_, 10 mM Ascorbic acid), overnight at 4°C, prior to processing for immunofluorescence staining.

### Organoid Culture

Isolated crypts were dissociated to single cells with TripLE express (Life Technologies) at 37°C, 5 minutes. Dissociated cells were filtered through a 40 μm cell strainer (Greiner) and suspended in growth factor reduced Matrigel (BD Biosciences). Organoids were grown in crypt media (Advanced DMEM/F12 (ADF) supplemented with 10 mM HEPES, 2 mM Glutamax, 1 mM N-Acetylcysteine, N2 (Gemini), B27 (Life technologies), Penicillin/Streptomycin (Sigma) supplemented with growth factors – ENR media (EGF (50 ng/ml, Invitrogen), Noggin (100 ng/ml, eBioscience) and RSpondin conditioned media produced from stably transfected L-cells (1:4). Chiron99021 (3 μM), Valproic acid (1 mM, Invitrogen) and Y27632 (10 μM, eBiosciences) were added to the culture for the first 48 hours. Organoids were passaged every 3-5 days by mechanically disrupting Matrigel and by washing and pipetting in ADF. Dissociated crypts were re-suspended in fresh Matrigel and grown in crypt media supplemented with growth factors.

For small molecule treatments, primary intestinal epithelial cells were cultured in ENR-CVY (ENR plus Chiron99021, Valproic acid and Y27632) for 3 Days, and then organoids were sub-cultured in ENR for two further days prior to the start of the experiment. Organoids were then treated with the stated small molecules for the indicated time periods. For induction of Unlicensed-G_1_, organoids were treated with Gefitinib (5 μM) coupled with removal of EGF from the crypt media. For re-activation/chase period, the media was removed and fresh growth factors added. All growth factors and inhibitors were replenished every 2 days throughout the experiment.

### Flow cytometry and cell sorting

Intestinal crypts were isolated and dissociated to single cells as described above. Isolated cells were filtered through 40 μm cell strainers (Greiner). For organoids, following one PBS wash, organoids were dissociated to single cells by incubation in TripLE express for 15 minutes at room temperature followed by manual disruption by pippetting. Cells were then extracted with CSK buffer for 20 minutes on ice, followed by fixation in 0.5% PFA (pH7.40, 15 minutes, room temperature). Cells were then washed once in 1% BSA and permeabilized with ice-cold 70% EtOH, 10 minutes. Cells were then washed in 1% BSA and re-suspended with primary antibodies (Mcm2, 1:500; GFP, 1:500; Ki67, 1:200) diluted in Working buffer (overnight, 4°C). Following two washes in working buffer, cells were re-suspended in secondary antibodies goat anti-mouse or anti-rabbit (Alexafluor647, 1:500 (Molecular Probes); Alexafluor488-Ki67, 1:400 (Clone SolA15, BD Biosciences), diluted in working buffer (1 hour, room temperature). After two washes in 1% BSA, cells were suspended in working buffer containing 15 μg/ml DAPI. Samples were analysed on a FACS Canto (BD Biosciences).

For cell sorting, cells were isolated from Lgr5-GFP mice as described above by treatment with TripLE express for 15 minutes, 37°C followed by filtration through 40 μm filters (Greiner). Cells were sorted in ADF supplemented with 1% FBS and DAPI (15 μg/ml). Sorting was performed using an Influx™ Cell sorter (BD biosciences). Cells were checked post-sort to ensure sample purity by re-examining Lgr5 expression in the sorted gates.

### Microscopy and Image analysis

Samples were imaged using a Zeiss LSM 710 microscope using a 40X LD Pan-Neofluar objective lens and immersion oil with a refractive index of 1.514. Z-stacks were acquired at optimal section intervals between 0.3 and 0.8 μm. For quantification, images were acquired at 16 bit-depth.

Image processing and analysis were performed using Imaris (Bitplane). Images of individual crypts were manually cropped, ensuring that an individual crypt was the only region of interest. All nuclei were detected in individual crypts using automated thresholding in Imaris using the measurement point function, set to detect nuclei at an estimated size of 3.5 μm. Missed or incorrectly assigned nuclei were manually identified. This function produced measurement points that segmented the specific region at the corresponding co-ordinate of the measurement point. Mean intensities for different channels were calculated per spot. This equates to the intensity at the centre region of each nucleus. A reference nucleus at the crypt base was used to define the crypt base position. The Euclidean distance to this point was measured and defined as the distance to the crypt base. Multiple images were analysed using the same workflow and the analysed files collated. For vibratome sections, a plane was manually defined running through to the muscle layer beneath the epithelium. The smallest distance to this surface was defined for segmented nuclei. For nuclear volume estimation, nuclei were manually segmented in 3D using the manual segmentation tools within Imaris (S1 Figure).

### Flow cytometry analysis

Flow cytometry data was analysed using FlowJo (Treestar) using a standardised gating strategy (S1 Figure). Briefly, cells were identified using FSC and SSC. Following doublet discrimination, gates were set using appropriate controls lacking conjugated secondary antibodies and without primary antibodies. Mcm2 negative gates were set by secondary only controls in conjunction with the Mcm2 intensity of G_2_ cells. G_1_ cells were discriminated based on the maximal DNA-bound Mcm2 intensity prior S-phase, and by DAPI intensity.

### Computational Modelling

A deterministic computer model for licensing in G1 was written in the Swift programming language using the Xcode 9 development environment. The model assumes that there is a minimum G1 period that is required for cells to grow to a critical size before they can enter S phase. Licensing can take place during this period at a constant rate. A licensing rate of 1 means that cells will be fully licensed in exactly the minimum G1 period. It is also assumed that cells have a robust ‘licensing checkpoint’ (Shreeram et al., 2002, Blow and Gillespie, 2008) so that they cannot enter S phase until origins have been fully licensed. An optional ‘unlicensed G1 period’ occurs at the start of G1, during which time no licensing takes place. In the simulation, cells enter G1, wait for any optional ‘unlicensed G1 period’, then start to license origins at a fixed rate; cells exit G1 and modelling ceases only when they have become maximally licensed and also the minimum G1 period has elapsed. The model divides the minimum G1 period into 10,000 equal steps and records the degree of licensing at the end of each step in the ‘Licensing Array’. The contents of the Licensing Array are distributed into 101 different frequency bins, ranging from 0% (no licensing) to 100% (maximal licensing). To model the background signal recorded by flow cytometry, all the licensing values are increased by 5%. To model flow cytometry measurement error, the frequency array is smoothed by starting at the smallest bin and pushing 80% of the cell counts into the next largest bin, and then starting at the largest bin pushing 80% of the cell counts into the next smallest bin.

~~~
//
// JBLicensingRateModel.swift version 1.0
//
// Created by Julian Blow on 05/10/2017.
import Foundation
class JBLicensingRateModel {
 let licensingRate: Double // rate of licensing in proportion to minimum G1 length
 let proportionUnlicensedG1: Double // proportion of minimum G1 before licensing occurs
 let timeStepsForMinimumG1: Int
 let licensingPerStep: Double
 let timeStepsBeforeLicensingStarts: Int
 let binsForFACS: Int
 let backgroundForFACS: Double
 let binSmoothingProportion: Double
 let totalNumberOfCells: Double
 let precisionCutOffForSmoothing: Double
 let numberOfTimepointsOutput: Int

 init (licensingRate: Double, proportionUnlicensedG1: Double) {
  self.licensingRate = licensingRate
  self.proportionUnlicensedG1 = proportionUnlicensedG1
  timeStepsForMinimumG1 = 10000
  licensingPerStep = (licensingRate / Double(timeStepsForMinimumG1)) * 100
  timeStepsBeforeLicensingStarts = Int(proportionUnlicensedG1 * Double(timeStepsForMinimumG1))
  binsForFACS = 100
  backgroundForFACS = 5
  binSmoothingProportion = 0.8
  totalNumberOfCells = 10000.0
  precisionCutOffForSmoothing = totalNumberOfCells / 10000 // cutoff for the smallest number of cells we continue to smooth for the far right of the array
  numberOfTimepointsOutput = 100
 }

 func run () -> ([Double], [Double]) {
  var licensingTimepoints: [Double]
  if licensingRate > 0 { licensingTimepoints = simulateLicensing() }
  else { licensingTimepoints = Array(repeating: 0.0, count: timeStepsForMinimumG1) } // if licensing can never occur
  let FACSOutputExact = simulateFACSExact(licensingTimepoints: licensingTimepoints)
  let FACSOutput = smoothFACSOutput(FACSOutputExact: FACSOutputExact)
  var simplifiedLicensingTimepoints = [Double]()
  for index in stride(from: 0, to: licensingTimepoints.count, by: licensingTimepoints.count/numberOfTimepointsOutput) {
   simplifiedLicensingTimepoints.append(licensingTimepoints[index])
  }
  return (FACSOutput, simplifiedLicensingTimepoints)
 }

 func simulateLicensing() -> [Double] {
 // simulates the degree of MCM loading given the licensingRate and proportionUnlicensed
 // returns amount of MCM loaded (percent max) at different time points
  var amountOfLicensing = 0.0
  var licensingArray = Array(repeating: 0.0, count: timeStepsBeforeLicensingStarts)
  var timeStep = timeStepsBeforeLicensingStarts

  repeat {
   amountOfLicensing += licensingPerStep
   if amountOfLicensing > 100 { amountOfLicensing = 100 }
   licensingArray.append(amountOfLicensing)
   timeStep += 1
  } while (amountOfLicensing > 100) || (timeStep < timeStepsForMinimumGI)
  return licensingArray
 }

 func simulateFACSExact (licensingTimepoints: [Double]) -> [Double] {
 // takes an array amount of MCM loaded at different time points as created by simulateLicensing()
 // turns this into a simulated output from a FACS (flow cytometry) analysis with totalNumberOfCells
  var maxMCMValue = licensingTimepoints.max()! // can be >100 because of smoothing
  if maxMCMValue == 0 { maxMCMValue = 100 } // if no licensing occurred
  var FACSOutputExact = Array(repeating: 0.0, count: binsForFACS+1) // binsForFACS+1 because we 0 and 100 are both possible
  let numberOfCellsPerTimepoint = totalNumberOfCells / Double(licensingTimepoints.count)   // so we get totalNumberOfCells events
  for anMCMContent in licensingTimepoints {
   let binIndex = Int(anMCMContent * Double(binsForFACS) / maxMCMValue)
   assert(binIndex <= binsForFACS, “Cannot have an MCM content greater than 100, the bin number”)
   if binIndex>=0 { FACSOutputExact[binIndex] += numberOfCellsPerTimepoint }    // add cells to bin
  }

  return FACSOutputExact
 }
 func smoothFACSOutput (FACSOutputExact: [Double]) -< [Double] {
  // simulates the error in FACS data by:
  // first adding a background of backgroundForFACS to all samples
  // then pushing binSmoothingProportion of each bin into the higher bin, when the value ” cutOffForDither
  // then pushing binSmoothingProportion of each bin into the lower bin
  let backgroundBins = Array(repeating: 0.0, count: Int(backgroundForFACS))
  var FACSOutput = backgroundBins + FACSOutputExact // add the background of empty pbins
  var binIndex = 0
  var amountToShift = 0.0
  repeat {   // expand rightwards by dithering
   amountToShift = FACSOutput[binIndex] * binSmoothingProportion
   FACSOutput[binIndex] −= amountToShift
   if binIndex+1 < FACSOutput.count { FACSOutput[binIndex+1] += amountToShift }
   else { FACSOutput.append(amountToShift) }
   binIndex += 1
   } while (amountToShift >= precisionCutOffForSmoothing) || (binIndex < FACSOutputExact.count + Int(backgroundForFACS))
  binIndex = FACSOutput.count-1
  while binIndex >= 0 {   // expand leftwards by dithering
   amountToShift = FACSOutput[binIndex] * binSmoothingProportion
   FACSOutput[binIndex] −= amountToShift
   if binIndex > 0 { FACSOutput[binIndex-1] += amountToShift }
   binIndex −= 1
  }

  return FACSOutput
 }
}
~~~

## RESULTS

### Mcm2 expression declines along the crypt-villus axis

Because of their abundance, strong conservation and association with the core DNA replication process, the presence of MCM2-7 proteins is commonly used to establish proliferative capacity in tissues, similar to Ki67 or PCNA (Gonzalez et al., 2005, Jurikova et al., 2016, Stoeber et al., 2001, Williams et al., 1998). Usually, terminally differentiated cells in mammalian tissues do not contain MCM2-7 (Stoeber et al., 2001, Todorov et al., 1998, Eward et al., 2004). To establish the overall MCM2-7 protein abundance along intestinal crypts, we first examined the expression of MCM2-7 proteins in adult murine small-intestinal epithelium using high-resolution immunofluorescence microscopy. We focused on Mcm2 as a surrogate for all the members of the MCM2-7 complex based on their similar function and localisation. However, we repeated a subset of the experiments using an antibody to Mcm4, which is less effective in detecting endogenous proteins. Nonetheless, in all cases, the results were identical.

Consistent with previous reports, Mcm2 was highly expressed in both murine and human intestinal epithelium. Mcm2 was highly expressed in intestinal crypts (Figure 1A) and declined gradually along the crypt-villus axis (Figure 1B), but persisted in a few cells in the villus compartment (Figure 1D). Mcm2 was nuclear in interphase cells but cytoplasmic during mitosis (Figure 1C). Although the majority of intestinal crypt cells expressed Mcm2, at the crypt base, Mcm2(+) and Mcm2(-) cells were interspersed (Figure 1A, D), consistent with previous reports (Pruitt et al., 2010). This pattern is reminiscent of the alternating arrangement of Lgr5(+) stem cells and Paneth cells at the crypt base (Barker et al., 2007). Lgr5(+) stem cells express Ki67 and are continually proliferative whereas Paneth cells are fully differentiated and are Ki67(-) (Basak et al., 2014). As expected, Mcm2 was expressed in all Lgr5(+) stem cells and there was a strong correlation between Mcm2 and Lgr5 expression (Figure 1E). This is consistent with the idea that Lgr5^Hi^ stem cells are the main proliferative stem cells in the intestinal crypt. Staining with Ulex Europaeus Agglutinin I (UEA), demonstrated that most Mcm2(-) cells in the crypt base are UEA(+) Paneth cells (Figure 1F).

**Figure 1.**
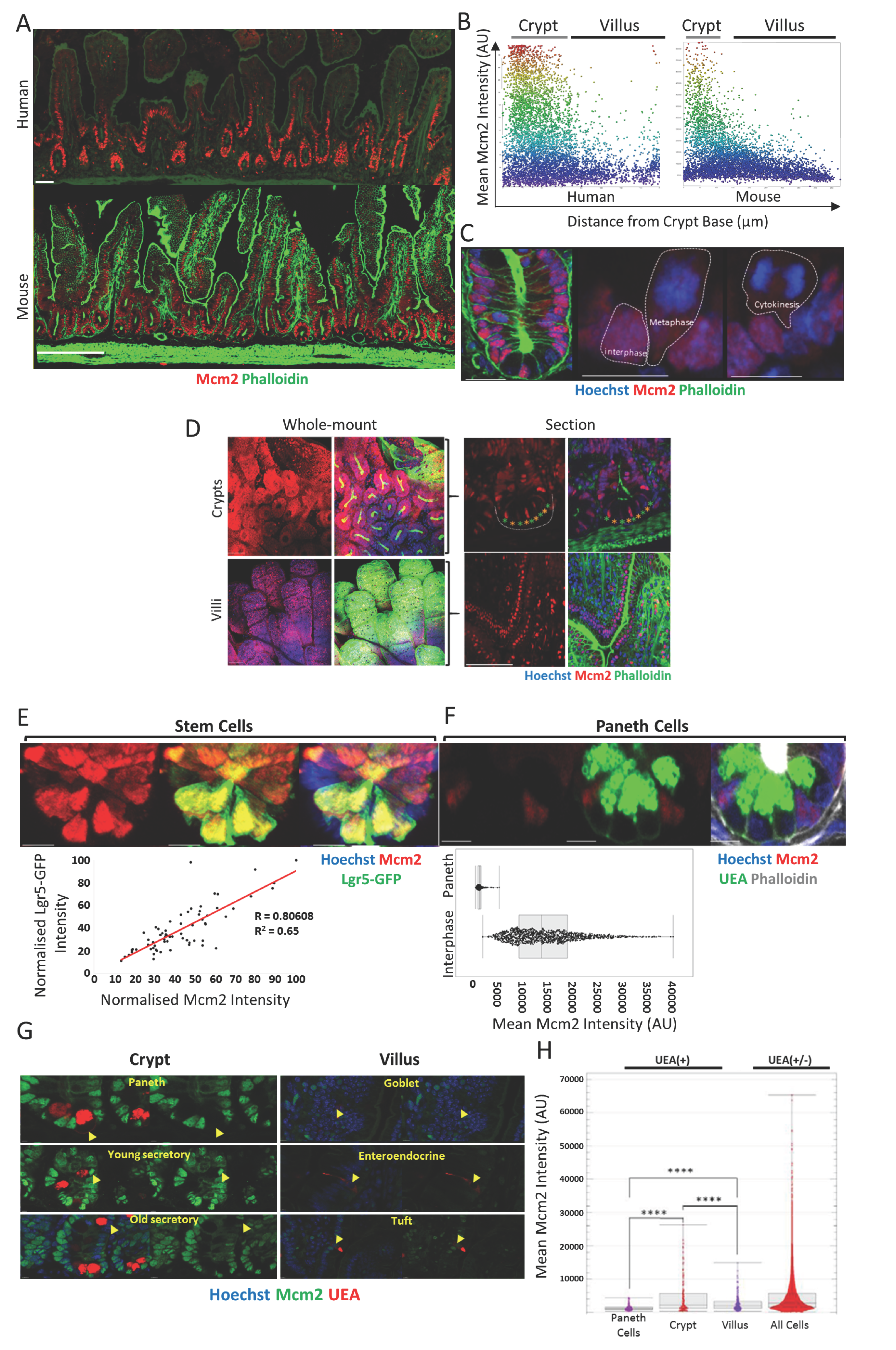
Mcm2 is expressed ubiquitously along the crypt-villus axis and declines slowly as cells differentiate. (**A**) Sections of normal human (top panel) and mouse (bottom panel) small-intestine were stained with Phalloidin (Green) and an antibody against Mcm2 (Red). Scale bars: 200 μm.
(**B**) Mean Mcm2 intensities for segmented nuclei were plotted along the crypt-villus axis for mouse and human tissue. Location of the crypt and villus domains is indicated.
(**C**) An intestinal crypt stained with Hoechst (Blue), Phalloidin (Green) and an antibody against Mcm2 (Red). Individual cells in interphase and mitosis (metaphase and cytokinesis) are outlined by dashed white lines.
(**D**) Maximum intensity projections of whole-mount intestinal tissue revealing intestinal crypts and villi (left panels). Individual X-Y sections are also shown to reveal the epithelium (right panels). Tissue was stained with Phalloidin (Green) and Hoechst (Blue) and an antibody against Mcm2 (Red). The alternating pattern of Mcm2(+) (Green stars) and Mcm2(-) (Orange stars) in the crypt base is highlighted)
(**E**) Images of Lgr5-GFP stem cells (Green) (top panel) co-stained with a Mcm2 antibody (Red). The correlation (Pearson’s correlation R=0.81, p<0.0001) between mean Mcm2 and Lgr5-GFP intensities for 69 Lgr5-GFP(+) cells normalised to the maximum intensity for an individual crypt is shown.
(**F**) Images of UEA(+) Paneth Cells (top panels) co-stained with an Mcm2 antibody (Red) and UEA (Green). Mean Mcm2 intensity for segmented nuclei of UEA(+) Paneth cells was compared with interphase cells (Right panels).
(**G**) Mcm2 (Green) and UEA (Red) expression in subsets of UEA(+) cells in crypt and villus domains. UEA(+) cells at the crypt base represent Paneth cells.
(**H**) Quantification of mean Mcm2 intensity in individual UEA(+) cell populations. UEA(+) cells in the crypt base (Paneth cells, N=224), in the upper crypt compartment (Crypt, N = 132) and in the villus compartment (Villus, N = 225) were identified manually, and the nuclear Mcm2 intensity was determined for individual cells (All cells, N = 33,736). There was a significant difference between UEA(+) cells in the crypt and villus compartments (T test, p<0.0001)

Normally, MCM2-7 expression is lost in terminally differentiated cells (Eward et al., 2004, Stoeber et al., 2001, Williams et al., 1998, Williams et al., 2004). The loss of expression has been suggested as a major contributor to the proliferation-differentiated switch *in vivo.* To test this idea, we measured the Mcm2 content of young and mature secretory cells in intestinal crypts and villi (Figure 1G, H). There was differential expression of Mcm2 in distinct secretory lineages. Many mature secretory cells including Paneth, Goblet and enteroendocrine cells were Mcm2(-), consistent with their differentiation status and long life-span in the epithelium (van der Flier and Clevers, 2009). We detected a small number of UEA(+) Mcm2(+) cells in intestinal crypts (Figure 1G). Assuming that Mcm2 expression declines slowly after terminal differentiation, the presence of Mcm2 in UEA(+) secretory cells could reflect their immaturity. Consistently, Mcm2 expression in UEA(+) cells in crypts was significantly higher than in villi (Figure 1H), supporting the idea that MCM2-7 are gradually lost upon terminal differentiation. Since MCM2-7 are highly abundant and have a long (>24 hour) half-life (Musahl et al., 1998), it likely that after cells differentiate, their MCM2-7 content declines at a slow rate, explaining why Mcm2 persists in the villus compartment.

### Visualisation of DNA replication licensing *In vivo*

MCM2-7 exist in three states: as hexamers free in the nucleoplasm, as double hexamers bound to DNA during late mitosis, G1 and S phase, or as CMG complexes at replication forks during S phase (Evrin et al., 2009, Gambus et al., 2011, Remus et al., 2009). To distinguish between DNA-bound and soluble forms, we developed a protocol involving a brief extraction of isolated crypts with non-ionic detergent to remove soluble MCM2-7. The remaining Mcm2 should mark cells whose origins are licensed for replication. Extraction did not visibly affect intestinal crypt integrity but made them more opaque compared to unextracted tissue (Figure 2A). The majority of cells in unextracted crypts were Mcm2(+) (Figure 2B, ‘Total’) similar to tissue sections and mirroring the ubiquitous expression of Ki67 along the crypt axis. After extraction, the majority of the Mcm2 content in cells was lost (Figure 2B, ‘Licensed’), with only 10-30% of cells maintaining high levels of Mcm2 (Figure 2C). After extraction, Mcm2(+) was not present in mitotic cells expressing phosphorylated histone H3, confirming the extraction procedure successfully removed non-DNA bound MCM2-7 proteins (Figure 2D).

**Figure 2.**
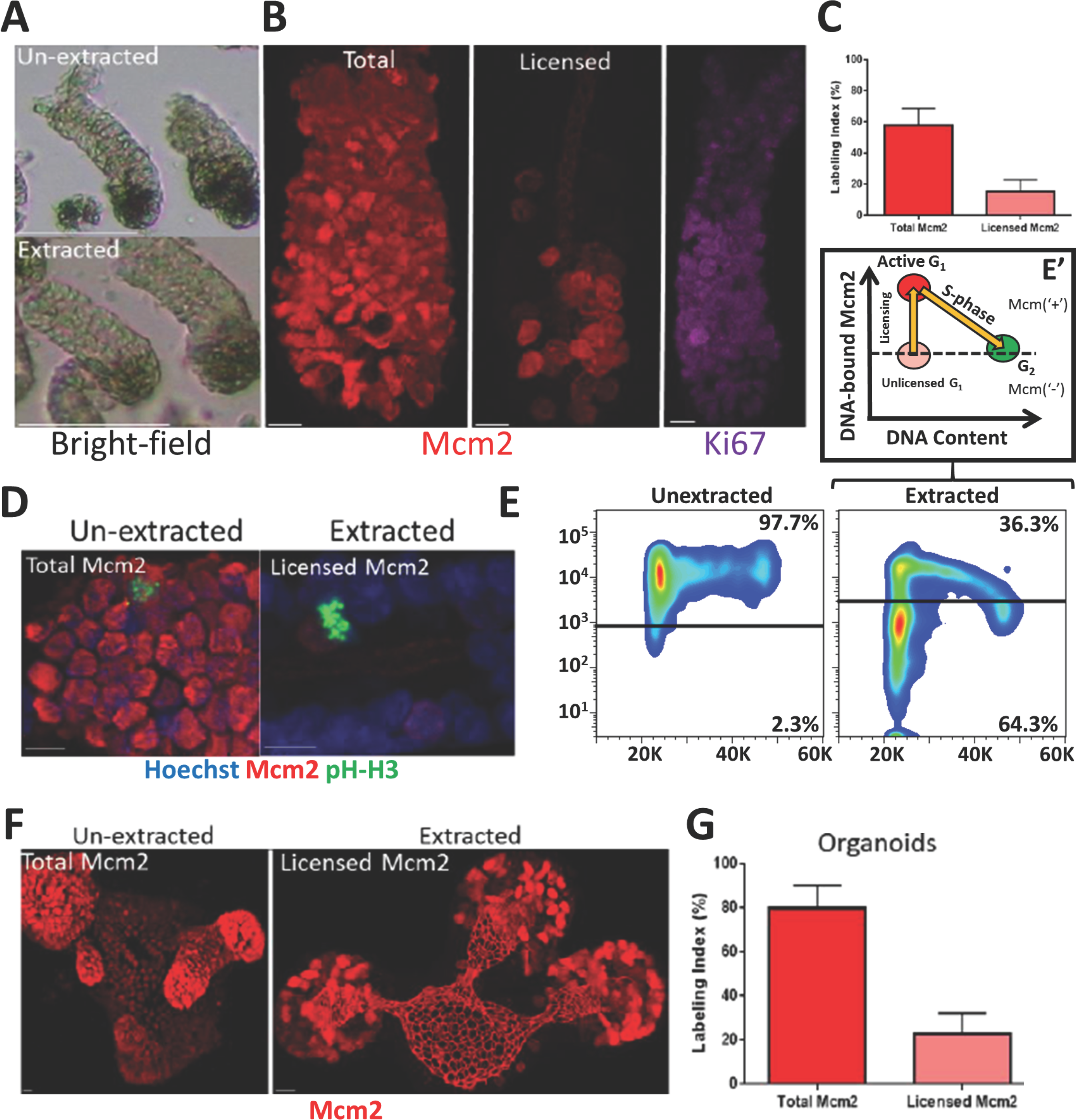
Visualising Mcm2 licensing in intestinal crypts. (**A**) Representative bright-field images of extracted and unextracted isolated intestinal crypts. Scale bar: 90 μm
(**B**) Representative images of isolated crypts stained with antibodies against Mcm2 (Red) or Ki67 (Purple). Scale bar: 10 μm
(**C**) The Mcm2 labelling index for unextracted and extracted crypts is significantly different (Mean +/-SEM, N=10 crypts (T test, p<0.0001).
(**D**) Representative intestinal crypts stained with Hoechst (Blue) and antibodies against Mcm2 (Red) and phospho-histone H3 (pH-H3) (Green).
(**E**) Representative flow cytometry profiles for extracted and unextracted isolated crypt epithelial cells showing Mcm2 vs. DNA content.
(**E’**) Suggested model of the licensing profile shown in panel E. Deeply quiescent cells, do not express Mcm2 and have a no detectable Mcm2 signal. Cells expressing soluble Mcm2 (Unlicensed G_1_) show a similar Mcm2 signal to G_2_ cells. After a proliferative fate decision has been made, origins become licensed and cells commit to S-phase entry. Cells enter S-phase after maximal origin licensing (Active-G_1_). During S-phase, Mcms are then displaced from DNA during replication.
(**F**) Representative images of extracted and unextracted intestinal organoids stained with an antibody against Mcm2 (Red).
(**G**) The Mcm2 labelling index for unextracted and extracted organoids. Data is displayed as Mean +/-SEM, N = 3 organoids and shows a significant difference (T test, p<0.0001).

We used flow cytometry to measure MCM2-7 content more directly and further confirm the effectiveness of the extraction procedure. Whereas the majority of isolated epithelial cells expressed Mcm2 that persisted throughout the cell-cycle, extraction revealed a distinct profile of Mcm-containing cells in crypts (Figure 2E). These profiles are consistent with those reported for cultured cell lines (Friedrich et al., 2005, Haland et al., 2015, Moreno et al., 2016, Matson et al., 2017). Mcm2 is present throughout the cell cycle (Figure 2E, ‘Unextracted’), but extraction shows that it binds to DNA throughout G_1_, reaching a maximum level before cells enter S-phase, and is subsequently displaced from DNA during S phase. This behaviour, which matches the known cell cycle behaviour of MCM2-7, confirms the efficiency of our extraction protocol. An antibody against Mcm4 produced similar results (data not shown). We observed that the majority of cells with a G1 DNA content appeared to be unlicensed, having a DNA-bound Mcm2 content similar to G_2_/M cells (Figure 2E, E’). This is substantially different from typical profiles observed in cultured cells lines in which most G_1_ cells are fully licensed (Friedrich et al., 2005, Haland et al., 2015, Moreno et al., 2016, Matson et al., 2017). Similar results were observed in cells isolated from intestinal organoids (Figure 2F, G).

### Licensing status and cell-cycle progression along the crypt-villus axis

Cell-cycle dynamics of intestinal stem and progenitor cells are highly heterogeneous (Pruitt et al., 2010). The majority of Lgr5(+) stem cells are considered to be continually proliferative, but with a much longer cell-cycle than transit-amplifying progenitor cells, which are most commonly found in S-phase (Schepers et al., 2011). To investigate proliferative fate decisions of intestinal epithelial cells, we used our MCM2-7 extraction in crypts where S phase cells were labelled in vivo with the nucleoside analogue EdU. We then used image analysis software to correlate Mcm2 content with cell-cycle stage along the crypt-villus axis (S1 Figure A-F). This allowed quantification of licensing in relation to the cell-cycle and 3D spatial information.

Figure 3A shows tissue labelled in vivo with a 1 hour EdU pulse followed by extraction of soluble MCM2-7. As expected, the majority of cells in the transit-amplifying compartment were labelled with EdU suggesting that most cells were in S-phase, consistent with early studies using BrdU and [^3^H]-thymidine labelling (Chwalinski and Potten, 1987). The patterns of replication foci were consistent with the reported S-phase replication timing programme (Rhind and G_1_lbert, 2013). Typically, all licensed cells had intense nuclear Mcm2 staining. Some cells completely lacked Mcm2 and EdU labelling, suggesting they were in either G_0_, very early G_1_ or in G_2_. Some cells were labelled with both Mcm2 and EdU. These double-labelled cells typically showed patterns of EdU labelling consistent with early to mid S phase and Mcm2 labelling of DNA compartments expected to replicate later in S phase. This relationship has been observed in tissue culture cells (Krude et al., 1996) and is consistent with the idea that DNA-bound MCM2-7 are displaced from chromosomal domains as replication is completed. Cells with late S-phase patterns of EdU labelling had little or no detectable Mcm2, consistent with the displacement of the majority of MCM2-7 by the end of S phase. We also measured nuclear volume, which increases during S phase and G_2_. This showed that nuclear volume increased up to two-fold in cells classified as S-phase and Late-S/G_2_ by Mcm2 and EdU staining (Figure 3B). This confirms our cell-cycle assignment and also suggests that most Mcm2(-) cells are in G_0_ or G_1_, rather than in G_2_.

**Figure 3.**
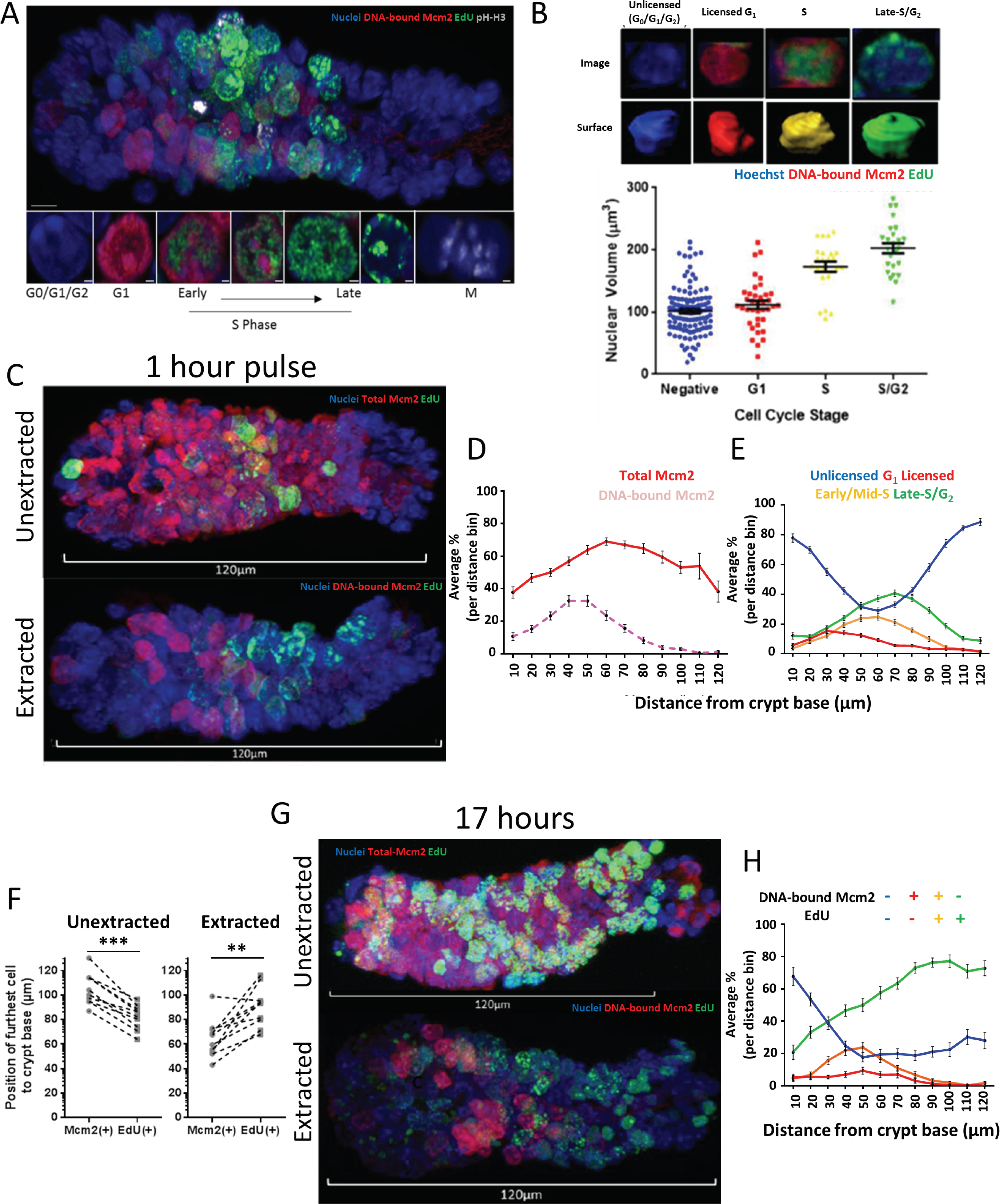
The licensing state defines distinct proliferative zones in intestinal crypts. (**A**) Representative image of an extracted intestinal crypt isolated after a 1 hour EdU pulse *in vivo* (EdU, Green) and stained with Hoechst (Blue) and antibodies against Mcm2 (Red) and pH-H3 (White). Co-staining shows distinct cell-cycle phases (bottom panels): Licensed cells committed in G_1_ (Mcm2(+), EdU(-)); Early (Mcm2(+), EdU(+)) to Late (Mcm2(-), EdU(+)) S-phase), and mitotic cells (pH-H3(+)). Negative cells represent deeply quiescent (G_0_), terminally differentiated cells or cells in G_1_, which have not made a proliferative fate decision, remaining unlicensed. The crypt base is to the left of the displayed image.
(**B**) Nuclear volume was estimated in cells at the distinct cell-cycle phases identified previously: Negative (G_0_/G_1_/G_2_: N =115), G_1_ Licensed (Mcm(+), EdU(-): N=38), S-phase (Mcm(+), EdU(+): N=24) and Late-S/G_2_ (Mcm(-), EdU(+): N=26). Top Panels show representative examples of each cell-cycle phase and the associated 3D rendered nuclei. There was a significant difference in the size of Licensed G_1_, S and Late-S/G_2_ nuclei (T test, p<0.0001).
(**C**) Representative images of intestinal crypts isolated after a 1 hour EdU pulse (Green) *in vivo.* Displayed are 3D projections of extracted and unextracted crypts stained with Hoechst (Blue) and an antibody against Mcm2 (Red). The crypt base is to the left of the displayed image.
(**D**) Comparison between cells expressing Mcm2 protein and DNA-bound Mcm2 along the crypt-villus axis between unextracted (N=101 crypts) and extracted (N=109 crypts) (taken from 3 mice). Data is displayed as the mean *%* of cells per set distance bin.
(**E**) All cells were divided into 4 distinct groups based on Mcm2 and EdU intensities. These groups represent distinctive cell-cycle phases as defined by their total (unextracted N=101 crypts) or licensed (extracted N=109 crypts) Mcm2 content: **Extracted:** 1) **Unlicensed** (Mcm2(-), EdU(-)), 2) **G_1_ licensed** (Mcm2(+), EdU(-)), 3) **Early/Mid S-phase** (Mcm2(+), EdU(+)) and 4) Late-S/G_2_ (Mcm2(-), EdU(+)). The data is represented as the population mean of the total cells per distance bin, Mean +/-SEM.
(**F**) The distance of the most distal Mcm2(+) and EdU(+) cells to the crypt base was compared in extracted and unextracted crypts. Data was scored manually for 10 representative crypts per condition. Licensed cells (Mcm2(+)) were significantly closer to the crypt base than EdU(+) cells (T test, p=0.0015. Cells expressing Mcm2 protein extended significantly above the last EdU(+) cell (T test, p<0.0003)
(**G**) Representative images of crypts isolated 17 hours after administration of EdU (Green). 3D projections of extracted and unextracted crypts stained with Hoechst (Blue) and an antibody against Mcm2 (Red) are shown.
(**H**) Cells were divided into 4 distinct groups as in Panel **E** (N=51 Crypts).

The combination of concurrently labelling DNA-bound Mcm2 and EdU showed a clear correlation between cell position and cell-cycle stage. As noted previously, Mcm2 is expressed in cells throughout the crypt (Figure 3C, D). At the base of the crypt, unlicensed cells predominate (Figure 3C, E). At increasing distances from the crypt base there is a successive rise in licensed G_1_ cells, early/mid S phase cells and then late S/G_2_ cells. Further up the crypt, at the end of the TA compartment, these cell cycle stages decline in reverse order, until unlicensed cells again predominate. This suggests that there is a co-ordinated progression through the cell division cycle as cells enter, then leave, the TA compartment. This was also observed as a field effect with many neighbouring cells showing similar replication patterns (S2 Figure A, B; S1 Movie).

### Terminal differentiation is associated with a binary licensing decision

At the terminal boundary of the transit-amplifying compartment, the majority of cells were unlicensed and had no DNA-bound Mcm2 (Figure 3C, E). Similarly, there were no licensed G_1_ cells beyond the TA compartment as defined by incorporation of EdU (Figure 3F). However, total Mcm2 expression extended significantly beyond the last cells with DNA-bound Mcm2 or incorporated EdU (Figure 3C, D, F). The distribution of total Mcm2 expression corresponded to the zone where cells express Ki67 (S3 Figure). Although Mcm2 and Ki67 expression persists beyond the TA compartment, licensing does not occur in this area. This suggests that differentiation is not governed by a gradual reduction in total MCM2-7 levels, but is a binary decision and licensing is abolished immediately after the final mitosis preceding differentiation. To further examine this, we marked the terminally differentiated zone by a 1 hour EdU pulse, followed by a 16 hour chase (Figure 3G, H). After 16 hours, the majority of the distal end of the TA compartment became labelled with EdU. All labelled nuclei in this area were significantly smaller than EdU(+) cells at the proximal end of the TA compartment (data not shown), suggestive of their differentiation status. Importantly the EdU(+) differentiated cells at the distal end of the TA compartment lacked DNA-bound Mcm2, supporting our suggestion that licensing is inhibited immediately at terminal differentiation.

### The majority of Intestinal stem cells spend the majority of G_1_ in an unlicensed state

The majority of cells in the crypt base expressed Mcm2, consistent with the finding that all Lgr5(+) cells express Mcm2 but mature secretory cells, such as Paneth cells, do not (Figure 1D, E). Surprisingly, extraction revealed that only 7-15% of cells were licensed in the crypt base (Figure 3C, D), with most cells in an unlicensed state despite expressing Mcm2. The abundance of licensed cells peaked 40-60 μm away from the crypt base, corresponding to just above the +4/+5 cell position (Figure 3D, E). This suggests that the majority of stem cells remain unlicensed. In contrast, the majority TA cells appear to progress rapidly through the cell-cycle, as many more actively incorporate EdU (Figure 4A).

**Figure 4.**
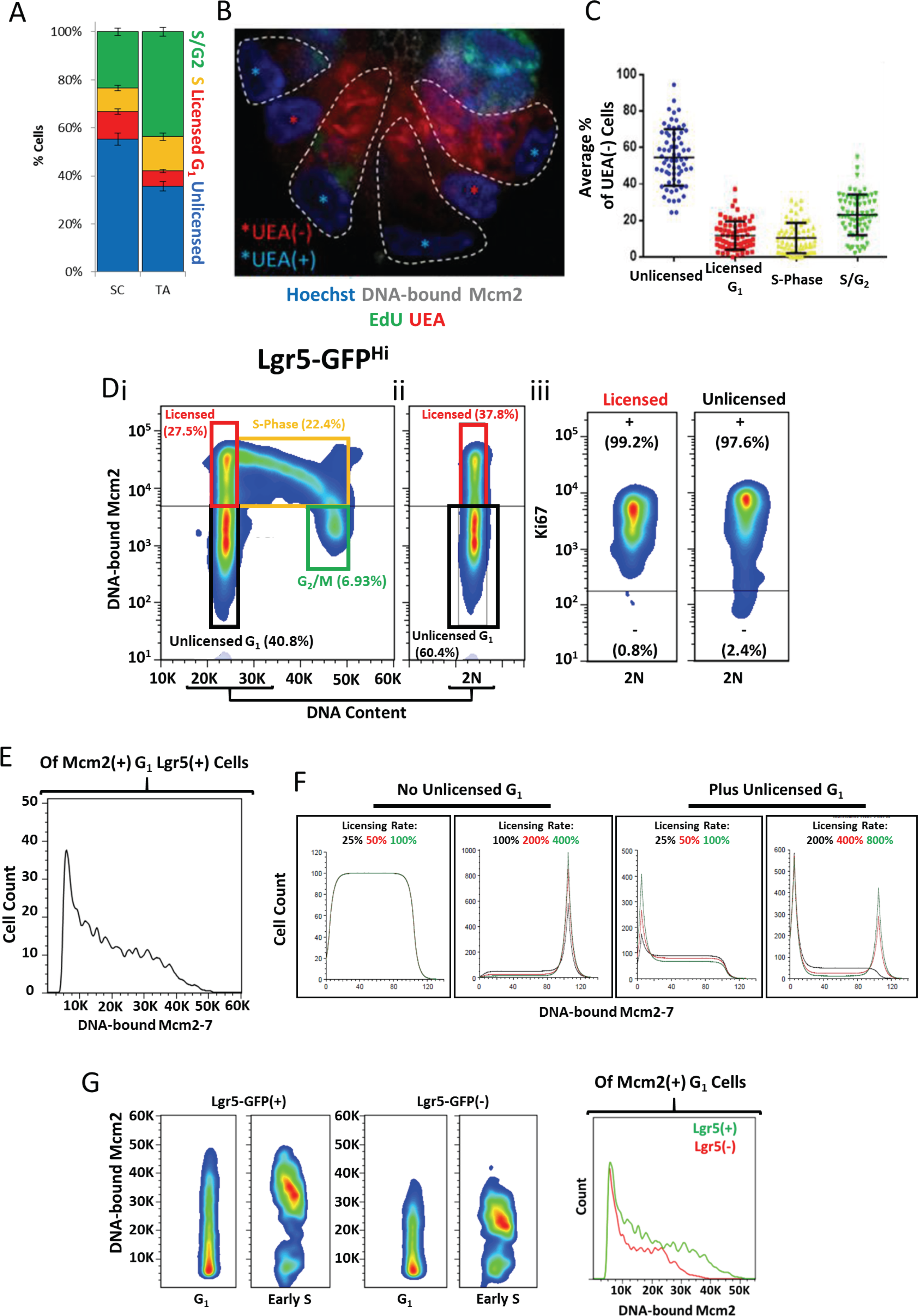
Intestinal stem cells reside in a paused unlicensed G_1_. (**A**) Quantification of cell cycle stages for cells in the stem cell (SC) compartment (<40 pm from the crypt base) and in the early transit-amplifying (TA) compartment (40-80 μm from the crypt base), as described in Figure 3E (**N=49 crypts**).
(**B**) Representative image of an extracted crypt base isolated 1 hour after a pulse of EdU (Green) and stained with Hoechst (Blue), UEA (Red) and antibodies against Mcm2 (White). Nuclear morphology and UEA signal were used to distinguish between UEA(+) Paneth Cells (outlined by dashed lines and nuclei marked with blue stars) and UEA(-) stem cells (situated between outlined Paneth cells and with nuclei marked by red stars).
(**C**) The average *%* of UEA(-) stem cells that fall into the previously defined cell-cycle bins: (Unlicensed; G_1_ licensed; S-phase; S/G_2_ phase; N=68 crypts).
(**D**) Representative flow cytometry profiles of (i) isolated Lgr5^Hi^ intestinal stem cells from 3 mice, showing DNA-bound Mcm2 and DNA content. The 2N (G_1_) cells of the same population are also shown (ii). The respective gated populations for Unlicensed G_1_ (Black), Licensed (Red), S-phase (Yellow) and G_2_/M (Green) are shown. The Ki67 content of the Licensed and Unlicensed G_1_ cells are also shown (iii).
(**E**) The frequency distribution of mean DNA-bound Mcm2 intensities for Mcm2(+) cells in G_1_ cells.
(**F**) Simulated ergodic rate analysis of origin licensing during G_1_. We simulated origin licensing dynamics during G_1_, varying the licensing rate and incorporating a significant paused period (Unlicensed G_1_). The displayed histograms show the frequency distribution of DNA-bound Mcm2 of G_1_ cells. Please refer to materials and methods for further information. However, an analogy to explain the model is as follows: Origin licensing through G_1_ can be thought of in terms of cars travelling along a long stretch of motorway: if cars enter the motorway at a fixed rate (i.e. unsynchronised cell cycles, cells entering G1) the density of cars at any one point is inversely proportional to the speed they are travelling at (‘speed’ in our case means the rate of licensing). We have added the concept of delays at the start and end, like tollbooths. In our model, there is a minimum drive time (the minimum length of G_1_ required for cells to grow to a critical size before they enter S phase (arbitrarily set at ‘100%’). If the speed of the cars means they don’t reach the end of the motorway before this time is up, cars maintain a constant speed along the entire road, and exit the motorway as soon as they reach the end. If cars drive faster and they reach the end before this time is up (>100%), they have to wait at the end of the motorway till the critical time is expired, resulting in a peak accumulation at the fully licensed point. Unlicensed G_1_ is an enforced time that cars have to wait once they enter the motorway before they are allowed to drive along it. This creates a peak on the left (unlicensed cells). If the cars then drive slowly, when they get to the end of the motorway, they exit immediately because the critical time has expired (centre panels), **but** if they drive fast enough they reach the end of the motorway before the critical time has expired. This results in a peak accumulation at both minimally and maximally licensed points.
(**G**) Comparison of DNA-bound Mcm2 content of Lgr5(+) and Lgr5(-) G_1_ cells and of cells in very early S-phase.

We next confirmed that the unlicensed cells at the crypt base were Lgr5(+) stem cells. As it is not possible to identify Lgr5 in these experiments, as it is extracted along with unbound Mcm2, we instead identified Paneth cells by UEA staining and considered all UEA(-) cells in the crypt base to stem cells (Figure 4B). >50% of the UEA(-) stem cells were in an unlicensed state and were not incorporating EdU (Figure 4C). Approximately 30-40% of all UEA(-) cells in the stem cell compartment were in an active phase of the cell cycle (licensed G_1_, S or G_2_; Figure 4C), corresponding to 5-6 stem cells out of the total 14 present (Snippert et al., 2010). This number is similar to the small number of proposed ‘working’ stem cells in the crypt base (Baker et al., 2014, Kozar et al., 2013). Unlicensed cells not incorporating EdU (i.e. unlabelled in this experiment) could theoretically be in either G_0_ or in G_2_. To distinguish between these possibilities we first isolated crypt cells from Lgr5-GFP mice and measured both GFP and DNA content. Both Lgr5(+) and Lgr5(-) cell populations had a similar cell-cycle profile with the majority of cells having 2N DNA content (S2 Figure C). We also examined the nuclear volume of cells at different positions along the crypt axis, after staining for EdU incorporation and DNA-bound Mcm2. The majority of unlicensed cells had a similar nuclear volume to that of fully licensed cells in G_1_ and not cells in Late-S/G_2_ phase (S2 Figure D). Together, these results suggest that, although they express abundant Mcm2, the majority of intestinal stem cells reside in an unlicensed G_1_ state.

To confirm this conclusion, we flow-sorted unextracted Lgr5-GFP(+) cells, extracted unbound MCM2-7 and stained them for Mcm2 and Ki67. Consistent with our previous results, most Lgr5(+) cells with a 2N DNA content had low levels of DNA-bound Mcm2, and were in an unlicensed state (Figure 4Di, ii). Importantly, both the licensed and unlicensed cells were Ki67(+) indicating that they had not withdrawn from the cell-cycle long-term (Figure 4Dii).

This unlicensed G_1_ state −2N DNA content, high Mcm2 expression but with low levels of DNA-bound Mcm2 - could be explained by two, slightly different, scenarios. 1) MCM2-7 are loaded on to DNA slowly in Lgr5(+) cells, thereby extending G_1_ (Schepers et al., 2011) (Dalton, 2015); or 2) Most Lgr5(+) cells enter G_1_ and remain in an unlicensed state but do not load MCM2-7 until an active decision is made to commit to cell cycle progression and activate the licensing system, at which time MCM2-7 proteins are rapidly loaded. In option 1) where licensing is slow, the presence of unlicensed cells simply reflects the increased time required to fully license origins, and different levels of Mcm2 loading should be equally distributed between G_1_ cells. In option 2), however, where there is an early stage of G1 where licensing does not occur, there should be a discrete peak of unlicensed cells with a G_1_ DNA content representing cells that have withdrawn from the cell cycle, and a lower frequency of G_1_ cells that have loaded different amounts of MCM2-7. To distinguish between these two possibilities, we utilized ergodic rate analysis. When examining the frequency distribution of DNA-bound Mcm2 in Lgr5(+) cells with 2N DNA content we found a discrete peak of unlicensed cells (Figure 4E) consistent with the second model. To confirm these observations, we performed computer modelling in which the licensing rate was varied in the presence or absence of an initial G_1_ period when licensing did not occur. (Figure 4F). With no unlicensed G1 period and low licensing rates (Figure 4F, left hand panels), G1 cells were equally distributed across the different degrees of licensing. With faster licensing rates and no unlicensed G1 period, a distinct peak of fully licensed G1 cells appeared, similar to what has been observed in cell lines. Only when an unlicensed G1 period was introduced did a distinct peak of unlicensed G1 cells appear (Figure 4F, left hand panels). Since most of these unlicensed Lgr5(+) cells express abundant Mcm2 (Figure 1D), this suggests that their G_1_ is characterised by a long unlicensed period. This may explain why the cell-cycle length of intestinal stem cells is significantly longer than transit-amplifying cells (Schepers et al., 2011).

To further confirm that many intestinal stem cells exist in an unlicensed G_1_ state, we examined the expression of Fluorescence ubiquitination cell-cycle indicator (FUCCI) reporters as an independent indicator of cell-cycle progression. We harvested intestinal tissue from Fucci2aR mice (Mort et al., 2014), and examined the expression of the G_1_ specific hCdt1(30/120) and S/G_2_/M specific hGeminin(1/110) reporters (S4 Figure A). As expected, many TA cells were hGeminin(1/110)(+). We also noticed that terminally differentiated cells, such as Paneth cells in the stem cell compartment, and cells at the tips of villi expressed high levels of hCdt1(30/120) (S4 Figure B) reflecting accumulation of the reporter in differentiated cells (Mort et al., 2014). However, we found that most Mcm2(+) stem cells in the crypt base expressed very low levels of the hCdt1(30/120) reporter, and only few cells expressed high levels (S4 Figure C, D). In contrast, the majority of TA cells were either hGeminin(1/110)^High^ or hCdt1(30/120)^low^, consistent with a short G_1_ phase. Together, this suggests that stem cells remain in a paused unlicensed G_1_ state where they do not rapidly accumulate hCdt1(30/120) unlike rapidly proliferating cells.

It has previously been reported that embryonic stem cells license more replication origins than neural stem/progenitor cells differentiated from them (Ge et al., 2015). To determine if adult stem and non-stem cells in intestinal crypts exhibit such differences, we compared the amount of DNA-bound Mcm2 in G_1_/G_0_ and early S phase Lgr5(+) cells with that in Lgr5(-) cells (Moreno et al., 2016). Although the majority of Lgr5(+) cells were unlicensed, when they entered S phase they had approximately twice as much DNA-bound Mcm2 compared to Lgr5(-) cells (Figure 4G). This is consistent with the idea that adult intestinal stem cells license more origins than TA cells and may represent a mechanism to protect genomic integrity.

### Intestinal label retaining cells are in a deep G_0_ state

Although the intestinal crypt base primarily consists of Lgr5(+) stem cells there is also a reserved pool of quiescent stem cells, often referred to as ‘+4 label retaining cells’ (LRCs) reflecting their position in the crypt base and their ability to retain nascent DNA labels (Potten et al., 2002). These cells are a rare subset of Lgr5(+) cells and are also secretory precursors (Buczacki et al., 2013). To further define the licensing status of these label-retaining intestinal stem cells, we identified UEA(-) LRCs by expressing H2B-GFP (which is incorporated into the chromatin of dividing cells) for 7 days and then chasing for a further 7 days (Buczacki et al., 2013, Roth et al., 2012). Labelled cells that did not divide during the 7-day chase period contain high levels of H2B-GFP (hence are LRCs), but cells that divided multiple times only have low levels. LRCs were then distinguished from Paneth cells based on UEA staining (Figure 5A). After induction, most cells in the epithelium expressed H2B-GFP (Figure 5Bi). After the 7D chase, H2B-GFP expression was restricted to cells near the villus tips, and cells at the crypt base (Figure 5Bii). We could successfully distinguish between Paneth cells and LRCs on the basis of UEA staining (Figure 5Bii). Unlike the majority of the Lgr5(+) cells, LRCs with high GFP-H2B did not express Mcm2 (Figure 5C). As expected, only non-LRC daughter cells with low levels of H2B-GFP had DNA-bound Mcm2 (Figure 5D). This shows that, LRC stem cells are in deep G_0_, unable to license because they do not express MCM2-7. In contrast, ‘active’ intestinal stem cells mostly reside in an unlicensed G_1_ state.

**Figure 5.**
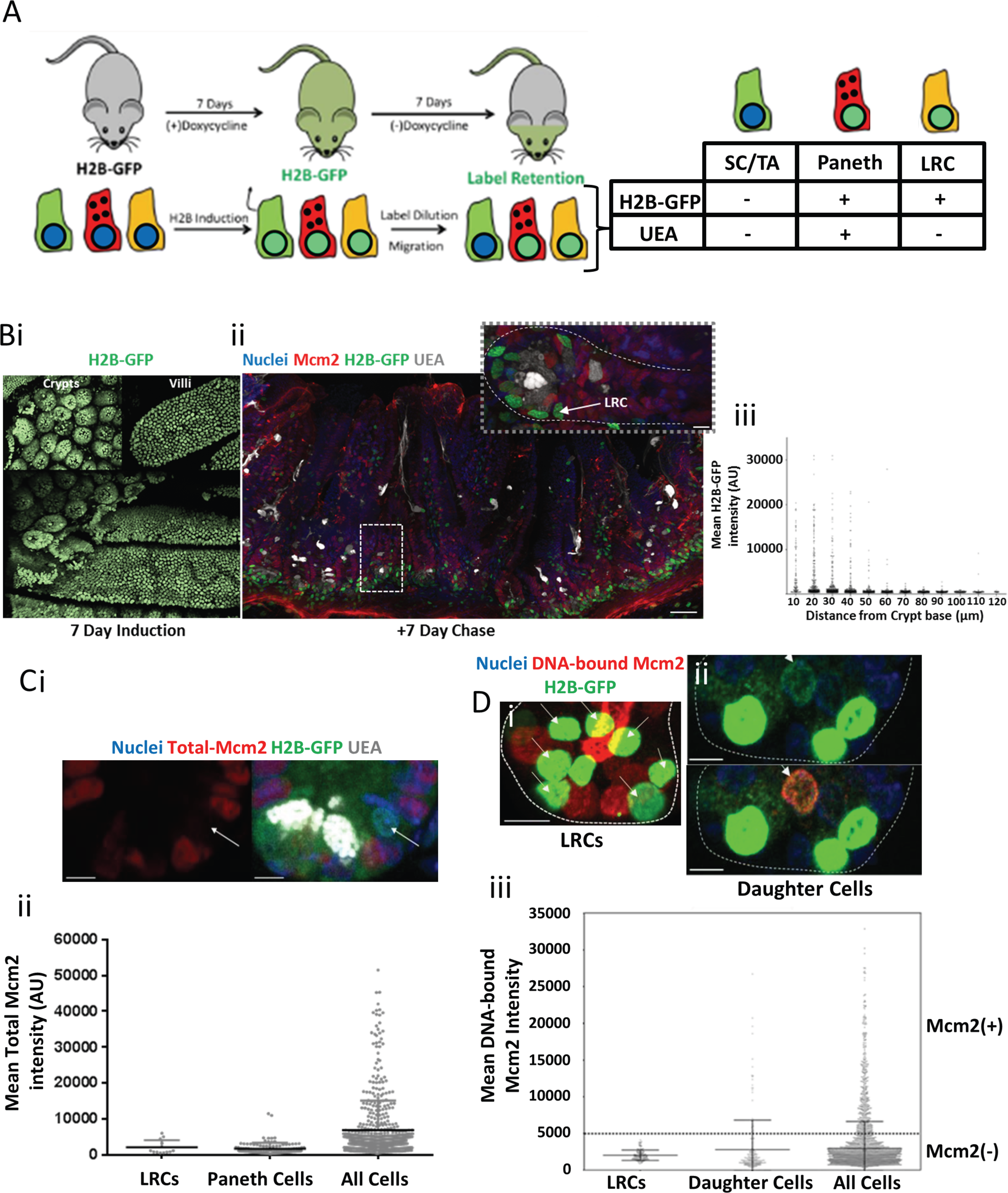
Label retaining cells are in a deep-Go state. (**A**) Labelling strategy. H2B-GFP expression was induced in all intestinal epithelial cells in H2B-GFP mice by administration of doxycycline for 7 days. After complete labelling, doxycycline was removed and mice rested for 7 days. During this chase period, the majority of H2B-GFP(+) cells are lost by label dilution due to cell division and upward migration. Both Paneth cells and +4 Label retaining cells are H2B-GFP(+) after the chase period. +4 Label retaining cells were distinguished from Paneth cells on the lack of UEA staining (Buczacki et al., 2013).
(**B**) Representative images of whole-mount sections of H2B-GFP expressing small-intestinal tissue after a 7-day labelling period (i). A vibratome section (ii) of intestinal tissue after a subsequent 7-day chase period, stained with Hoechst (nuclei), UEA (Grey), and an antibody against Mcm2 (Red). An enlarged image of the marked crypt is shown. A UEA(-) label retaining cell (LRC) is marked by the white arrow). Quantification of the mean GFP intensity for all cells along the crypt-villus axis after the 7 days chase period is shown (iii).
(**C**) A representative image (i) of a label-retaining cell in an intestinal crypts stained with Hoechst (Blue), UEA (White) and an antibody against Mcm2 (Red). The label retaining cell (White arrow) does not express Mcm2. The quantification of Mcm2 expression in label retaining cells (N=12), Paneth cells (N= 116) and all cells (N=543) is shown (iii).
(**D**) Representative images of extracted H2B-GFP crypts after the 7-Day chase period. H2B-GFP^Hi^ cells (bright Green) represent Paneth cells and +4 cells (i), and H2B-GFP^Low^ cells (Faint Green) represent daughter cells (white arrow) that have diluted H2B-GFP content as a result of cell division (ii). The quantification of DNA-bound Mcm2 in GFP^Hi^ label retaining (LR) and GFP^low^ daughter cells compared with the total cell population (all cells) is shown (iii).

### Licensing dynamics in intestinal organoids

Intestinal stem cells reside in a highly specialised niche at the base of crypts. It is therefore possible that this niche specifically allows stem cells to pause in an unlicensed G_1_ state where origin licensing is prevented until a further proliferative fate signal is received. To understand the contribution of the stem cell niche to the dynamics of entry and exit from the unlicensed G_1_ state, we used intestinal organoids, which allowed us to manipulate the stem cell niche/environment with small-molecules. Although the distribution of licensed cells was similar between the branches of intestinal organoids and crypts in tissue, there were considerably more cells with DNA-bound Mcm2 in the former (Figure 6A, Ciii). Importantly, cells in organoids showed a discrete peak of fully licensed G_1_ cells in addition to the cells in the unlicensed G_1_ state (Figure 6Bii). This suggests that the epithelium in organoids represents an accelerated state of self-renewal and may not fully recapitulate cell-cycle dynamics of intestinal epithelial cells *in vivo.*

**Figure 6.**
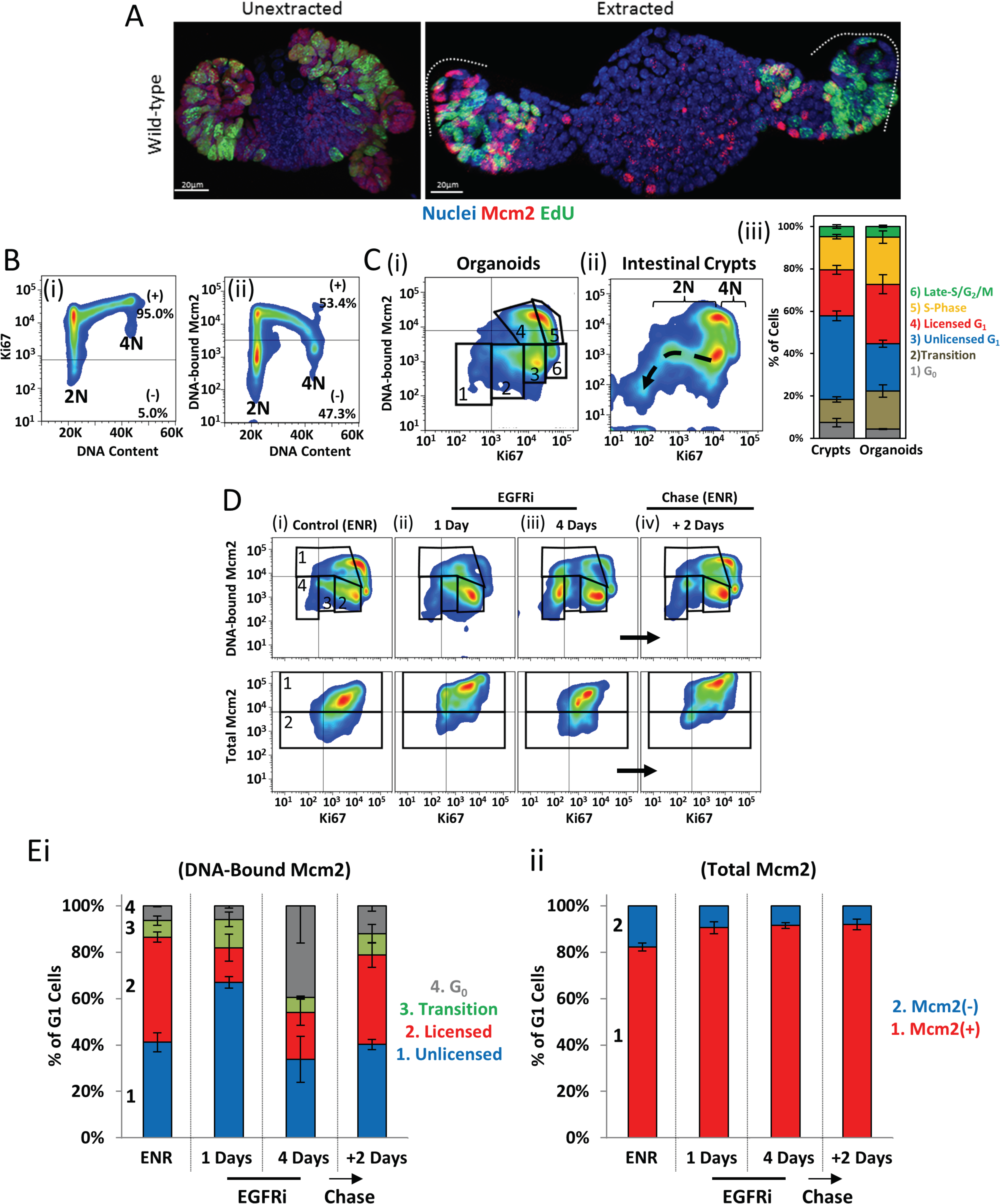
Licensing dynamics in intestinal organoids. (**A**) Representative image of an unextracted and extracted intestinal organoid stained with Hoechst (Blue), and an antibody against Mcm2 (Red) after a 1 hour EdU pulse (green).
(**B**) Representative flow cytometry profiles from isolated organoids cells showing Ki67 (i) or DNA-bound Mcm2 (ii) content vs DNA content.
(**C**) Representative flow cytometry profile from cells isolated from cultured organoids (i) or intestinal crypts (ii) showing DNA-bound Mcm2 vs Ki67 (**See also** S5 Figure A). Ki67 loss and subsequent loss of Mcm2 during differentiation is apparent, starting from Unlicensed G_1_ (dashed arrow). Quantification of discrete populations (gates shown in i) was performed: 1) G_0_, 2) Transition (G_0_0 Unlicensed G_1_), 3) Unlicensed G_1_, 4) Licensed G_1_, 5) S-Phase, 6) Late-S/G2/M. (N=3).
(**D**) Representative flow cytometry profiles of isolated organoid epithelial cells grown in ENR (control) and treated with the EGFR inhibitor Gefitinib, for the indicated times (i-iii). After 4 Days in Gefitinib, organoids were reactivated by removal of the Gefitinib and re-addition of fresh growth factors (ENR) for 2 days (iv). Displayed are profiles comparing DNA-bound Mcm2 vs Ki67 (Top panels) or total Mcm2 content (Bottom panels).
(**E**) The G_1_ cell populations as shown in (D) were quantified. DNA-bound Mcm2 profiles (i) show 1) Unlicensed G_1_, 2) Licensed G_1_, 3) Transition (G_0_↔ Unlicensed G_1_), 4) G_0_ populations. Total-Mcm2 profiles (ii) show Mcm2 expressing (Mcm2(+)) and non-expressing (Mcm2(-)) Cells. (N=3 organoids)

We designed an assay to robustly assess licensing dynamics during entry and exit from the unlicensed G_1_ state, and transition towards G_0_. Specifically, we used flow cytometry based quantification of DNA-bound Mcm2, Ki67 and DNA content to measure licensing dynamics during entry and exit from the unlicensed G_1_ state, and transition towards G_0_ (S5 figure A). Most cells in organoids express Ki67 and it increased during cell-cycle progression (Figure 6Bi). The DNA-bound Mcm2 profile was similar to that in isolated crypts (Figure 6Bii). Correlating Ki67 and DNA-bound Mcm2 produced a distinctive profile that is similar for isolated cells from organoids (Figure 6Ci) and intestinal crypts (Figure 6Cii). This profile reveals a population of cells that appear to lose Ki67 (dashed arrow in Figure 6Cii) and may represent cells decreasing in proliferative capacity and transitioning towards differentiation (S5 Figure A). Such a loss of proliferative capacity appeared to initiate in cells that express Ki67 but are unlicensed, i.e. cells in unlicensed G_1_. These data suggest that different stages of quiescence can exist that reflected by a spectrum of Ki67 and Mcm levels.

Stem-cell niche maintenance in organoids mainly depends on a combination of EGF, Wnt and Notch signalling (Sato et al., 2011). To identify the pathway that can modulate the unlicensed G_1_ state, we systematically treated organoids with a small molecule inhibitor of EGFR (Gefitinib), a Wnt agonist (Chir99021), and a Notch activator (Valproic acid). Short-term treatment with Gefitinib, which reduces MAP kinase activity and blocks DNA replication and cell division (Lynch et al., 2004), immediately caused cells to accumulate in the unlicensed G_1_ state, with a 2N DNA content but continuing to express Mcm2 and Ki67 (Figure 6D, E). Only prolonged EGFR inhibition (4 days) caused transition to an intermediate G_0_ state, with significantly reduced Ki67 expression but with total Mcm2 levels maintained (Figure 6D, E). Both states were reversed by removal of EGFRi and addition of fresh growth factors (Figure 6D, E). Previously it was shown that EGFR increases Lgr5 expression (Basak et al., 2017), suggesting that these transitional states (Unlicensed G_1_ and G_0_) are associated with ‘stemness’. Additionally, EGFRi appeared to potently kill transit-amplifying cells (S5 Figure B, C), leaving branches containing only stem cells (Basak et al., 2017). This suggests that both, the fully quiescent G_0_ and unlicensed G_1_ states, may provide protection to stem cells.

Treatment with the Wnt agonist Chir99021 did not appear to significantly affect licensing dynamics (S5 Figure C). Strikingly, treatment with the Valproic acid (a Notch activator) alone or in combination with Chir99021 significantly altered licensing profiles (S5 Figure D, F). The combination of Chir99021 and Valproic acid induces Lgr5 expression throughout the organoid epithelium (S5 Figure E) (Yin et al., 2014). Our data showed that this is associated with the appearance of a population of cells with low levels of Ki67 and intermediate levels of DNA-bound Mcm2 (S5 Figure D) similar to the intermediate, shallow G_0_ state induced by EGFRi. Surprisingly, we observed an arc of cells connecting this to the fully licensed state, suggesting that re-licensing of these cells occurs before they express high levels of Ki67 and that cells can reactivate licensing from a deeper state of G_0_ directly (S5 Figure D, F).

Inhibiting Notch signalling with DAPT, which induces terminal secretory cell differentiation (van Es et al., 2005), also significantly altered licensing dynamics and induced deep G_0_, with reduced Ki67 and loss of Mcm2 proteins (S5 Figure G). Together, these data suggest that both EGFR signalling and Notch signalling can significantly influence the licensing dynamics during the transition between quiescence and unlicensed G_1_.

### The Unlicensed-G_1_ state is lost in *Apc* mutant organoids

Many established cell lines appear to lack an unlicensed G_1_ state, instead licensing all origins immediately upon mitotic exit (Friedrich et al., 2005, Haland et al., 2015, Moreno et al., 2016, Matson et al., 2017). In addition, many of these cell-lines lack a functional licensing checkpoint (Shreeram et al., 2002, Feng et al., 2003, Liu et al., 2009, Nevis et al., 2009) that has been suggested to arrest cells in G_1_ by inactivation of the RB-E2F restriction point(Liu et al., 2009, Machida et al., 2005, Nevis et al., 2009, Shreeram et al., 2002, Teer et al., 2006). To understand the biological relevance of the unlicensed G_1_ state, we determined whether origin licensing dynamics during G_1_ are altered during the initial stages of tumorigenesis.

The first initiating mutations in colorectal cancer are usually in *Adenomatus polyposis coli* (Apc). Therefore, we investigated whether licensing dynamics are altered in *Apc* mutant intestinal epithelium. In Apc^Min/^+ epithelium *in vivo*, Mcm2 expression appeared normal in areas of normal histology but was greatly increased in polyps (Figure 7A). In isolated Apc^Min/+^ crypts, there was a slight increase in the size of the proliferative compartment similar to previous reports (Trani et al., 2014), and a slight increase in the number of EdU(+) cells in the stem cell compartment (Figure 7Bi, iii). However, in the majority of crypts there were no significant differences in the number or distribution of licenced cells in the stem cell compartment (Figure 7Bi, iii). However, we found a small number of crypts that were considerably larger. Strikingly, within these crypts, we noticed small ‘ribbons’ of EdU(+) cells that extended significantly into the transit-amplifying compartment (Figure 7Bii). We suspected that these ribbons reflected a clone of cells that had undergone loss of heterozygosity (LOH), and continually re-engage with the cell cycle. This in turn suggests that the LOH event that converts *Apc^Min/+^* to *Apc^Min/Min^* cells significantly alters cell-cycle dynamics.

**Figure 7.**
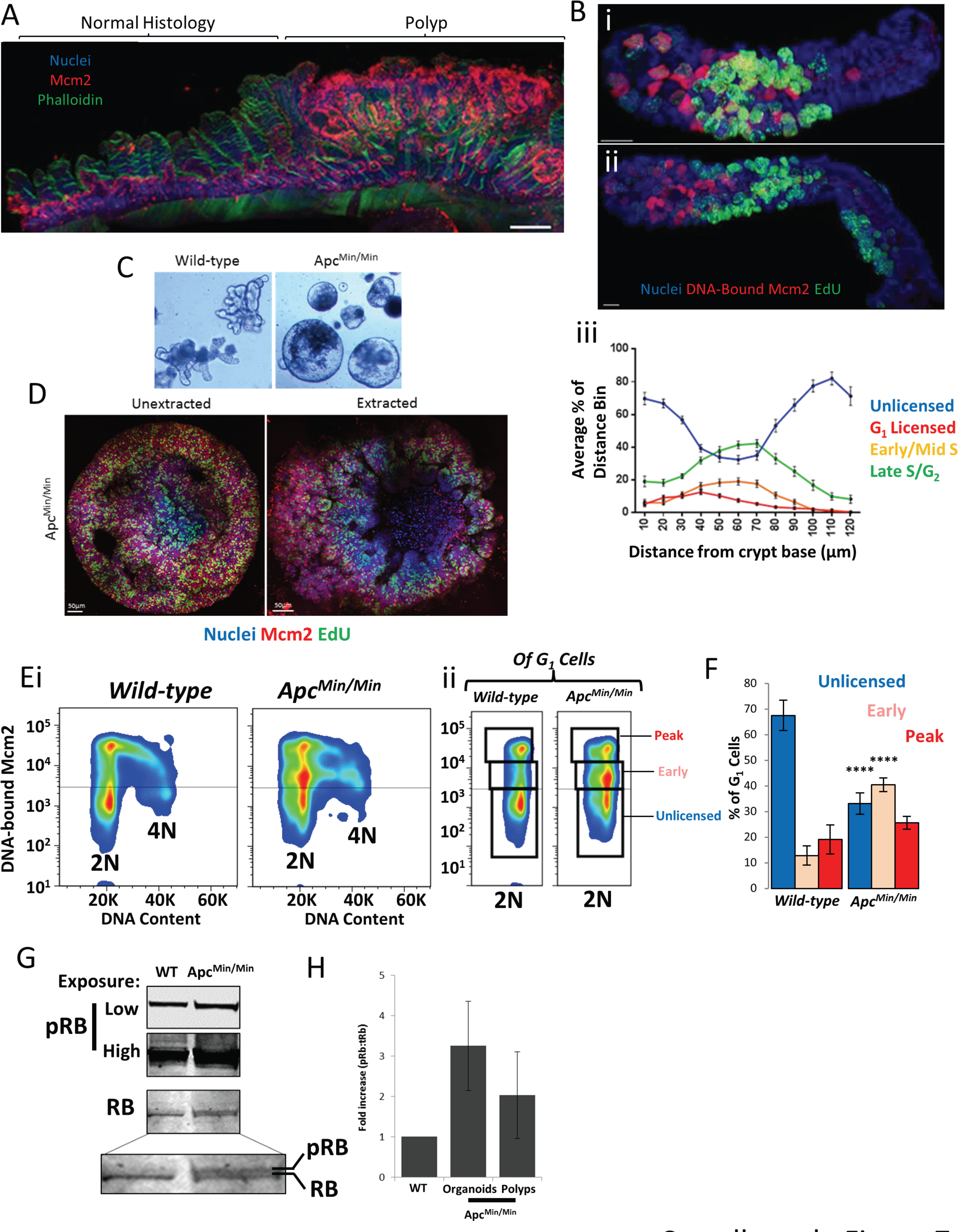
Unlicensed G_1_ is lost in *Apc* mutant epithelium. (**A**) A representative vibratome section of *Apc^Min/^+* intestinal epithelium stained with Hoechst (nuclei), Phalloidin and an antibody against Mcm2. Regions of normal histology and a region containing a polyp are highlighted.
(**B**) Representative image of an extracted *Apc^Min/^+* isolated crypt (i) stained with Hoechst (Nuclei) and an antibody against Mcm2 (Red), after a 1 hour EdU pulse (Green). A representative image of an abnormally elongated crypt is displayed (ii) showing a clonal ribbon of cells in the distal transit-amplifying compartment which are EdU(+). Quantification of cell-cycle stages across the crypt axis (Figure 3) is shown (iii).
(**C**) Representative bright field images of organoids cultured from wild-type and organoids from Apc^Min/^+ mice that have undergone loss of heterozygosity (Apc^Min/Min^).
(**D**) Representative images of unextracted and extracted *Apc^Min/Min^* organoids stained with Hoechst (nuclei), an antibody against Mcm2 (red), after a 1 hour EdU pulse.
(**E**) Representative flow cytometry profiles from cells from extracted wild-type and *Apc^Min/Min^* organoids showing DNA-bound Mcm2 vs DNA content (i). The 2N G_1_ cells of the shown profile are displayed (ii) showing Unlicensed G_1_, Early G_1_ and Peak (fully licensed) G_1_ populations.
(**F**) Quantification of the populations described in Panel Eii. Data is displayed as mean +/-SEM, N=3 organoids.
(**G**) Western blot of RB and pRB (low and high exposures) levels between wild-type and *Apc^Min/Min^* organoids. Two bands corresponding to hypo-and hyper-phospohorylated RB can be observed.
(**H**) Quantification of the fold increase of pRB:RB for Apc^Min/Min^ organoids (N=3) and polps isolated from Apc^Min/^+ mice (N=6), compared to *wild-type* organoids and tissue.

To determine whether LOH alters origin licensing dynamics by modifying G_1_, we compared wild-type and *Apc^Min/Min^* organoids (Figure 7C). *Apc^Min/Min^* organoids contained many more licensed cells that were distributed randomly throughout the organoid (Figure 7D), reminiscent of the altered distribution of Ki67 (+) cells in these organoids (Fatehullah et al., 2013). Strikingly, we found that licensing dynamics in the G_1_ phase of *Apc^Min/Min^* cells were significantly different and there was a significant loss of the unlicensed G_1_ population. (Figure 7E). Instead, there was a significant population of cells that appear to license immediately upon G_1_ entry and progressed into S-phase immediately after minimal licensing (Figure 7E, F). Interestingly, we still detected a population of cells that licensed as many origins as in WT organoids. These may be *Apc^Min/+^* cells that had not undergone LOH yet and thus maintained near-normal cell cycle dynamics.

A functional licensing checkpoint depends on an intact RB restriction point and we tested whether there the relative amounts of hypo-and hyper-phosphorylated RB varied between wild type and *Apc^Min/Min^* epithelia. In *wild-type* organoids, we noticed that the majority of RB appeared hypo-phosphorylated (Figure 7G). In contrast, in *Apc^Min/Min^* organoids, at least half of RB appeared hyper-phosphorylated, with at least 3-fold more phosphorylated RB. Together, this data is consistent with the idea that cells exist in an unlicensed G_1_ state prior to passage through the restriction point and activation of E2F-driven transcription. In contrast, *Apc* mutant cells appear to have lost normal restriction point control, so that they have constitutively hyper-phosphorylated RB, allowing them to completely bypass the unlicensed G1 state.

## Discussion

The cell-cycle of intestinal stem and transit-amplifying cells is poorly understood. By comparing the total and DNA-bound Mcm2 in intact intestinal crypts we provide new insights into how licensing and cell-cycle commitment are coupled in this tissue. We provide evidence that after their final mitosis, transit amplifying cells do not license their replication origins and immediately exit the cell cycle. We show that many Lgr5(+) stem cells spend the majority of G_1_ in an unlicensed state, continually expressing Mcm2 and Ki67. In unlicensed-G_1_, stem cells could be poised to respond to cues and progress past this restriction point to resume the cell division cycle.

Lgr5(+) stem cells have a cell-cycle length greater than transit-amplifying cells (Schepers et al., 2011). The biological relevance of this is currently unknown. The data presented here suggest a delay in origin licensing is a key feature of the prolonged cell-cycle of Lgr5(+) cells. Although ~80% of Lgr5(+) cells are thought to be continually proliferative and express high levels of both Ki67 (Basak et al., 2014) and Mcm2, we found that most Lgr5(+) cells reside in an unlicensed state, with 2N DNA content and Mcm2 not bound to DNA. Since the licensed state defines proliferative fate commitment (Blow and Hodgson, 2002), we suggest that these cells are temporarily paused in G_1_, continuing to express proliferative makers such as Ki67 and Mcm2, but without fully committing to the cell cycle (Figure 8). We show that the number of Lgr5(+) cells with DNA bound-Mcm2 was similar to the number of proposed ‘active’ stem cells determined in lineage tracing experiments (Baker et al., 2014, Kozar et al., 2013).

**Figure 8.**
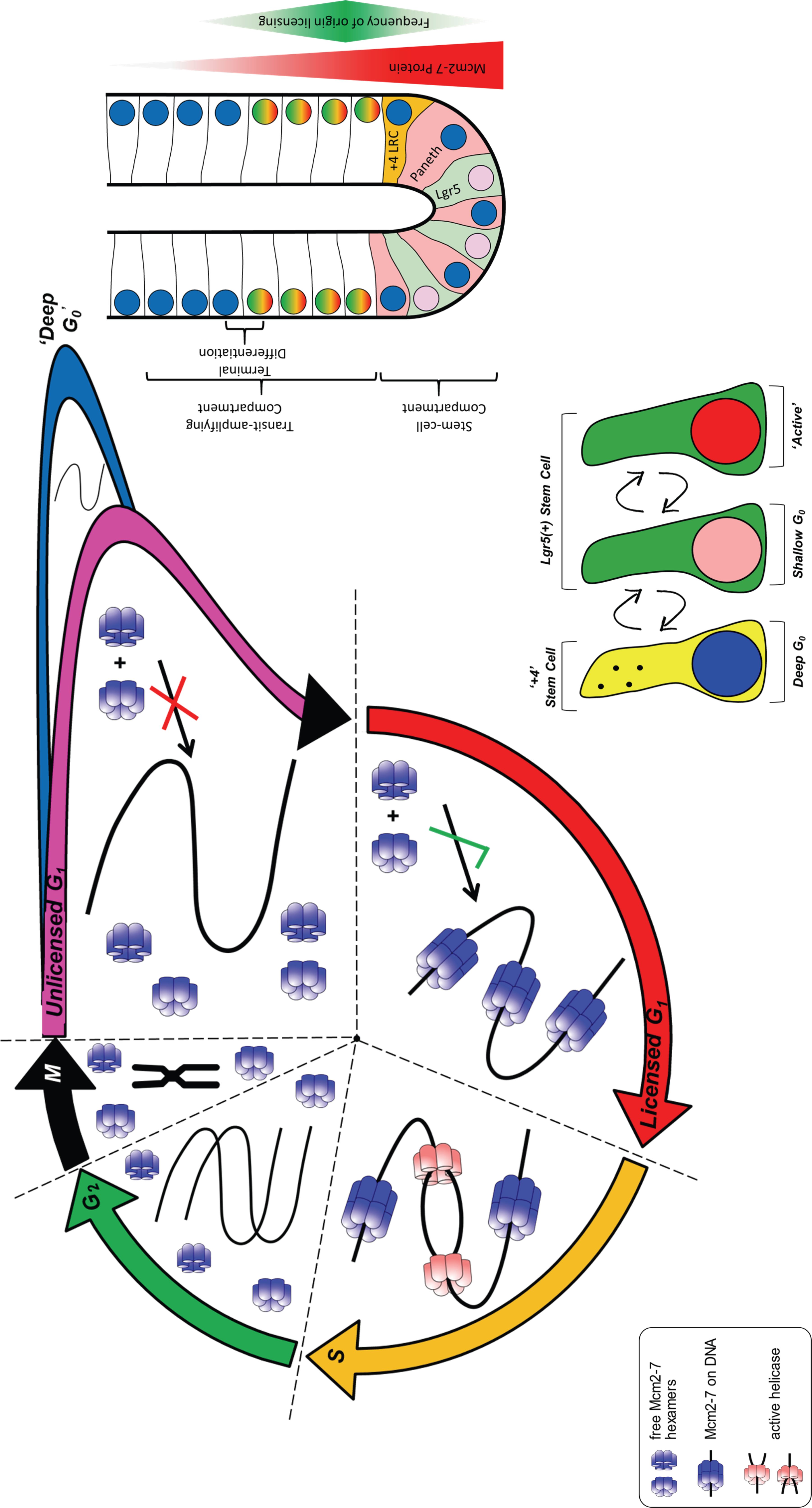
Model of Origin licensing dynamics in intestinal epithelial cells. In a normal cell-cycle, Mcms are expressed ubiquitously in all stages. The licensing of DNA with MCM2-7 occurs in late M and throughout G_1_, when a cell receives a stimulus to commit to the cell cycle. As DNA is replicated during S-phase, MCM2-7s are displaced from DNA and are prevented from relicensing in G_2_. During terminal differentiation, MCM2-7 are not actively transcribed and the proteins are gradually lost in post-mitotic cells. However, after the final mitotic division, cells make a binary decision never to license their DNA, even though the protein is still present. Mcm proteins then degrade slowly, where cells enter a terminally differentiated state (deep G_0_). Alternatively, cells can exit mitosis, not relicense their DNA but maintain proliferative markers and disengage from the cell cycle for some time (Unlicensed G_1_). Two major classes of intestinal stem cells exist: ‘Active’ stem cells, engaged with the cell-cycle, and reserve, quiescent Label Retaining Cells. Label retaining cells are in a state of ‘deep’ quiescence, and do not contain MCM2-7 because they have disengaged from the cell cycle for some time. In this study, we show that most ‘Active’ Lgr5(+) stem cells reside in an unlicensed state, but contain MCM2-7 proteins. These cells reside in an ‘Unlicensed-G_1_’ until they make a proliferative fate decision, enter the cell-cycle and license. This provides an explanation for the elongated cell-cycle of Intestinal stem cells: They reside in a partial resting state where they may be able to respond to niche cues to divide. This therefore may constitute a unique mechanism to control stem cell numbers.

Prolonged arrest may eventually result in degradation of MCM2-7 proteins and lead to induction of a state of deep quiescence (G_0_). Consistent with this idea, we observed that LRCs, thought to provide a reserve of quiescent stem cells, did not express Mcm2. The lack of Mcm2 expression may reflect that a significant period of time has passed since these cells divided. The delay in activating the licensing system may create a prolonged time-window for Lgr5(+) cells to receive and interpret environmental cues before deciding to commit to duplication, offering a means to control their number. It is likely that the majority of Lgr5(+) cells regularly resume their cell cycle, given their continual expression of proliferation markers (Basak et al., 2014). The identity and decisions of Lgr5(+) cells are governed by stochastic choices and the ability to pause briefly in G_1_ offers them unique flexibility in making these choices.

As expected, the transition from unlicensed G_1_ to licensed G_1_ seems dependent on EGFR signalling. However, other pathways responsible for stem-cell maintenance can also significantly cause the appearance of a unique population of unlicensed cells with distinct cell-cycle dynamics. Growing evidence suggests that intestinal stem cell fate is not governed by asymmetric segregation of fate determinants (Lopez-Garcia et al., 2010, Snippert et al., 2010, Steinhauser et al., 2012). Instead, factors operating in the stem cell niche, such as Wnt and Notch signalling affect stem cell fate decisions and also reduce the cycle rate of intestinal stem cells (Hirata et al., 2013). This is consistent with the idea that cell fate choices are affected by decreasing proliferation rate and increased G_0_/G_1_ length. Indeed, extending G_1_ in mouse and human embryonic stem cells can drive differentiation (Calder et al., 2013, Coronado et al., 2013). Similarly, long G_1_ phases are associated with the generation of fate-restricted progenitors during neurogenesis (Arai et al., 2011). An extended time window in the cell-cycle has been suggested to allow niche factors and/or fate determinants to accumulate to direct progenitor fate (Calegari and Huttner, 2003). In the case of adult intestinal stem cells, holding cells in G_1_ may allow an extended time for stem cell fate factors to act and maintain stem cell fate. In contrast, many embryonic stem cells licence rapidly and the cell cycle slows throughout differentiation (Matson et al., 2017).

However, like embryonic stem cells (Ge et al., 2015), intestinal stem cells appear to have licensed more origins than non-stem cells when they enter S phase. This may help ensure accurate and complete genome duplication in long-lived stem cells (Moreno et al., 2016). With a higher demand for licensed origins, intestinal stem cells may therefore more readily engage the licensing checkpoint that ensures that all origins are licensed before cells enter S phase (Alver et al., 2014, Liu et al., 2009, Shreeram et al., 2002). This additional demand for licensed origins in stem cells may also explain why crypts hypomorphic for Mcm2 have a stem-cell deficiency (Pruitt et al., 2007).

It is unclear how intestinal stem cells enter a significant unlicensed G_1_ state. The simplest explanation is that licensing factors such as Cdt1 or Cdc6 are not readily available in newborn stem cells, and their synthesis has to be stimulated by an upstream signal for fate commitment via activation of E2F-driven transcription. This is the situation after prolonged quiescence, which is accompanied by passive downregulation of licensing factors (Coller, 2007). In contrast, in continually dividing cells their levels are maintained. Consistent with this idea, licensing factors such as Cdc6, along with many cyclin-CDK complexes, are down regulated beyond the end of the TA zone (Frey et al., 2000) (Smartt et al., 2007). Cells without a functional restriction point, such as *Apc* mutant cells or most cancer cell lines, could immediately license their origins upon entering G_1_ and can progress into S-phase without sufficient origins being licensed. Interestingly, both transit-amplifying cells and highly proliferative *Apc* mutant cells are highly sensitive to replication inhibitors, such as Gefitinib (S5 Figure H). Wild-type stem cells survive this treatment, potentially by engaging the licensing checkpoint to reversibly stall in Unlicensed G_1_. This suggests that the unlicensed G_1_ can protect stem cells from replication inhibitors and offers a potentially selective means to kill highly proliferative cells, such as *Apc* mutant cells (Shreeram et al., 2002, Blow and G_1_llespie, 2008).

In summary, we demonstrate that the dynamics of the DNA replication licensing system provides a new way for measuring the proliferative fate of intestinal stem cells. We suggest a model for ‘working’ intestinal stem cells that spend a significant proportion of G_1_ in a unlicensed state until a proliferative fate decision is made. Correspondingly, exit from the cell-cycle in label retaining ‘+4’ cells leads to loss of proliferative capacity and loss of Mcm2 expression causing cells to enter a deeply G_0_ quiescent state (Figure 6). The unlicensed G_1_ state is lost in *Apc* mutant epithelia, which lack a functional Rb-restriction point. We suggest that the unlicensed G_1_ state serves stem cells in controlling their numbers by regulating the cell-cycle.

## Author contributions

T.D.C, J.J.B and I.N conceived and designed the study; T.D.C and I.P.N collected the data; Y.C assisted with organoid experiments; T.D.C performed the data analysis. J.J.B performed the modelling simulations. T.D.C, I.N and J.J.B wrote the manuscript.

## Acknowledgements

We would like to thank members of the Näthke and Blow laboratories for general assistance and helpful discussions; Dr Paul Appleton, Dr Graeme Ball and the Dundee Imaging and Tissue Imaging Facility for support with microscopy and image analysis. The imaging facility is funded by the Welcome Trust Technology Platform award (097945/B/11/Z) and Welcome Trust award (101468/Z/13/Z). We thank Dr Rosemary Clarke and the Dundee Flow Cytometry Facility for support with flow cytometry, cell sorting and analysis. We also thank Dr Richard Mort (University of Edinburgh) for Fucci2aR mouse tissue. This work was supported by a programme grant from Cancer Research UK to I.N (C430/A11243) and to J.J.B (C303/A14301), Wellcome Trust grant WT096598MA and an MRC studentship award to T.D.C.

---

**S1 Figure.**
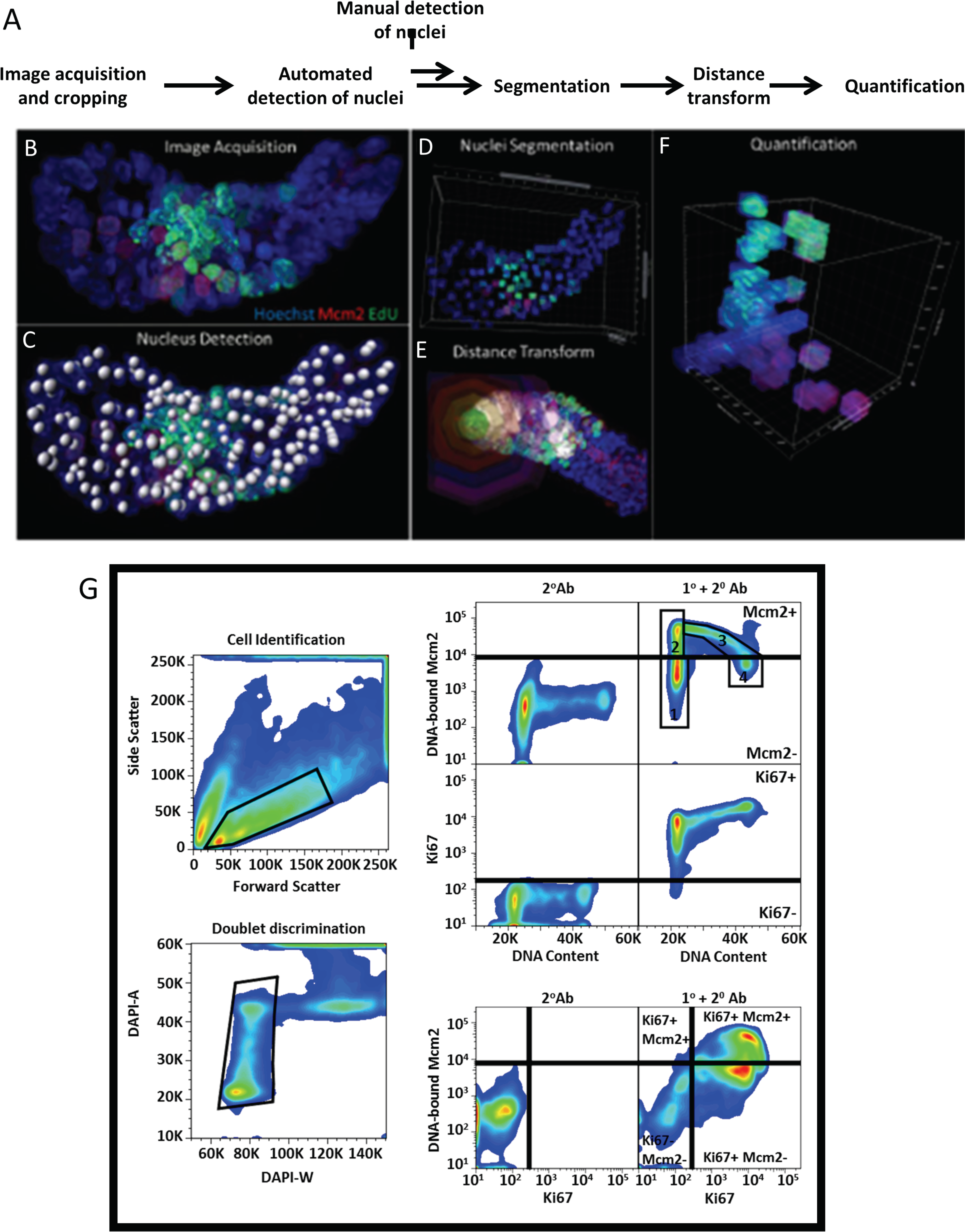
Image analysis. (**A**) Image analysis work-flow.
(**B**) Representative image of an extracted isolated crypt 1 hour after an EdU pulse (Green) stained with Hoechst (Blue) and an antibody against Mcm2 (Red).
(**C**) Detection of nuclei in the crypt in panel A in 3D. Nuclei were detected in 3D using segmentation tools in Imaris. Detection was validated visually or each individual crypt.
(**D**) Segmentation of the region of interest defined by nuclei detection in panel B.
(**E**) A distance transform was performed in Imaris to measure the distance of each nucleus to a reference nucleus at the crypt base. Visual representations of distances divided into different bins are displayed (Green, 0-10μm; Yellow, 10-20 μm; Red, 20-30μm; Blue, 30-40μm; Magenta, 40-50μm).
(**F**) Representative 3D quantification of the crypt in panel A shows the distance from the crypt base (X-axis), DNA-bound Mcm2 (Y-axis) and EdU incorporation (Z-axis).
(**G**) An overview of the gating strategy used for flow cytometry experiments. Intestinal epithelial cells were separated from debris on the basis of forward and side scatter. Single cells were subsequently distinguished from doublets based on pulse-width gating. Negative gates were set on control samples stained with secondary antibody alone. Unlicensed G_1_ (Population 1) and licensed-G_1_ (Population 2) were distinguished based on DNA-content and based on the threshold set by the negative control samples. S-phase cells (Population 3) can be seen as an arc of cells emanating from the fully licensed cell population, increasing in DNA content whilst losing DNA-bound Mcm2. G_2_/M cells (Population 4) can be seen as a population with 4N DNA content and lacking DNA-bound Mcm2.

**S2 Figure.**
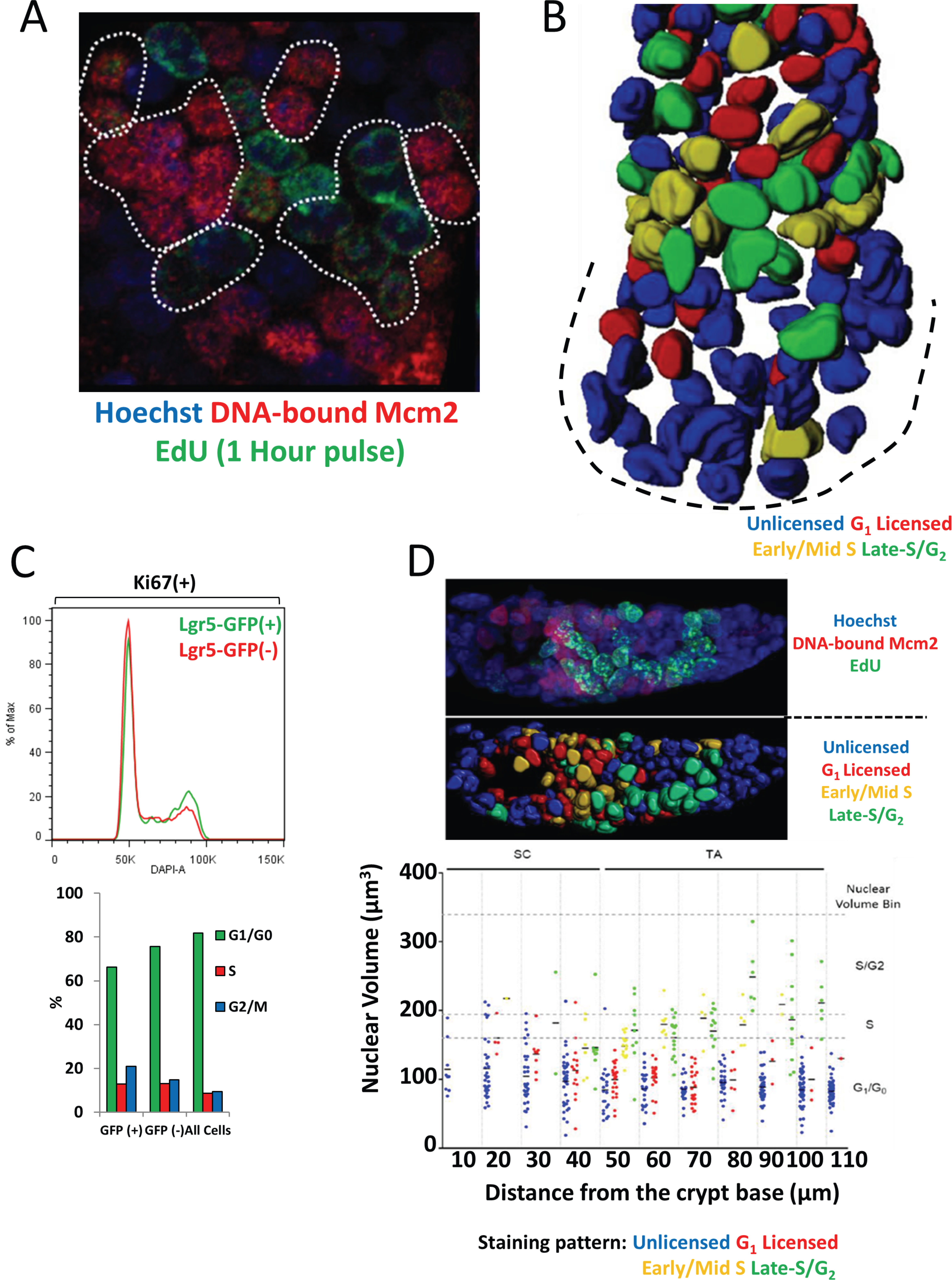
Clonal cell-cycle patterns in the intestinal epithelium. (**A**) Representative section through an extracted crypt after a 1 hour EdU pulse (Green) and stained with Hoechst (Blue) and an antibody against Mcm2 (Red). Discrimination of cell-cycle staging using DNA-bound Mcm2 and EdU incorporation patterns allows visualisation of clonal cell-cycle field effects revealing many neighbouring cells with similar DNA-bound Mcm2 and DNA replication patterns. These clones may represent lineages of from single cells that progress through the cell cycle at the same rate.
(**B**) Representative image of an isolated crypt in which surface rendering was performed on all nuclei and colour codes applied to reflect cell-cycle stage. Representative cell-cycle distributions for isolated Ki67(+), Lgr5(+) and Lgr5(-) intestinal epithelial cells are shown.
(**C**) Representative cell-cycle distributions for isolated Ki67(+), Lgr5(+) and Lgr5(-) intestinal epithelial cells. The average of each cell-cycle phase is displayed for duplicate isolations.
(**D**) Nuclear volumes were rendered for individual nuclei in whole intestinal crypts isolated 1 hour after labelling with EdU. Image shows nuclei (Blue), EdU (Green) and licensed Mcm2 (Red). Maximum intensity projections of the original image are displayed at the top and corresponding rendered nuclei at the bottom. Nuclear surfaces were colour-coded according to cell-cycle states: Blue, unlicensed; Red, Licensed G_1_; Yellow, S-phase; Green, Late-S/G_2_. Nuclear volumes were measured for all nuclei in representative crypts (N=3). Unlicensed (N=368); G_1_ (N=104); S-phase (N=41); Late-S/G_2_ (N=70) and the distance of cells from the crypt base were binned into 10μm intervals. Known parameters of the nuclear volume for known cell-cycle stages (Figure 3B) are overlaid.

**S3 Figure.**
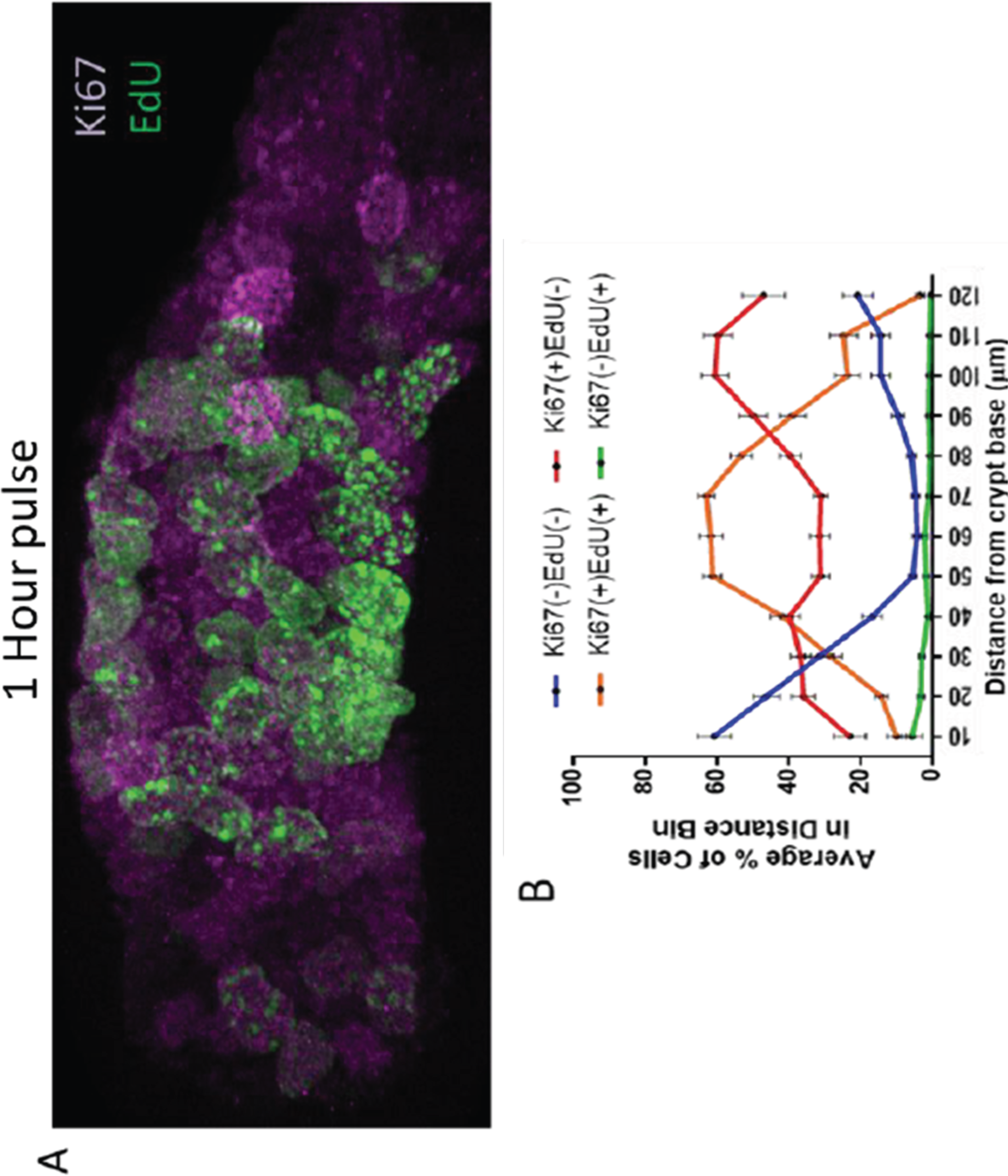
Ki67 expression along the crypt-villus axis. (**A**) A representative isolated crypt 1 hour after labelling with EdU (Green) and stained with an antibody against Ki67 (magenta)
(**B**) Quantification of the distribution of Ki67(+) cells along the crypt axis. Cells were binned into four groups: Negative (Ki67(-), EdU(-)); Ki67(+), EdU(-); Ki67(+), EdU(+) and Ki67(-), (EdU(+) and by their distance from the crypt base. Data is displayed as the average percentage of a particular cell subtype, per distance bin. Data is displayed as Mean +/-SEM. Data from 75 crypts is displayed, N = 14,264 cells.

**S4 Figure.**
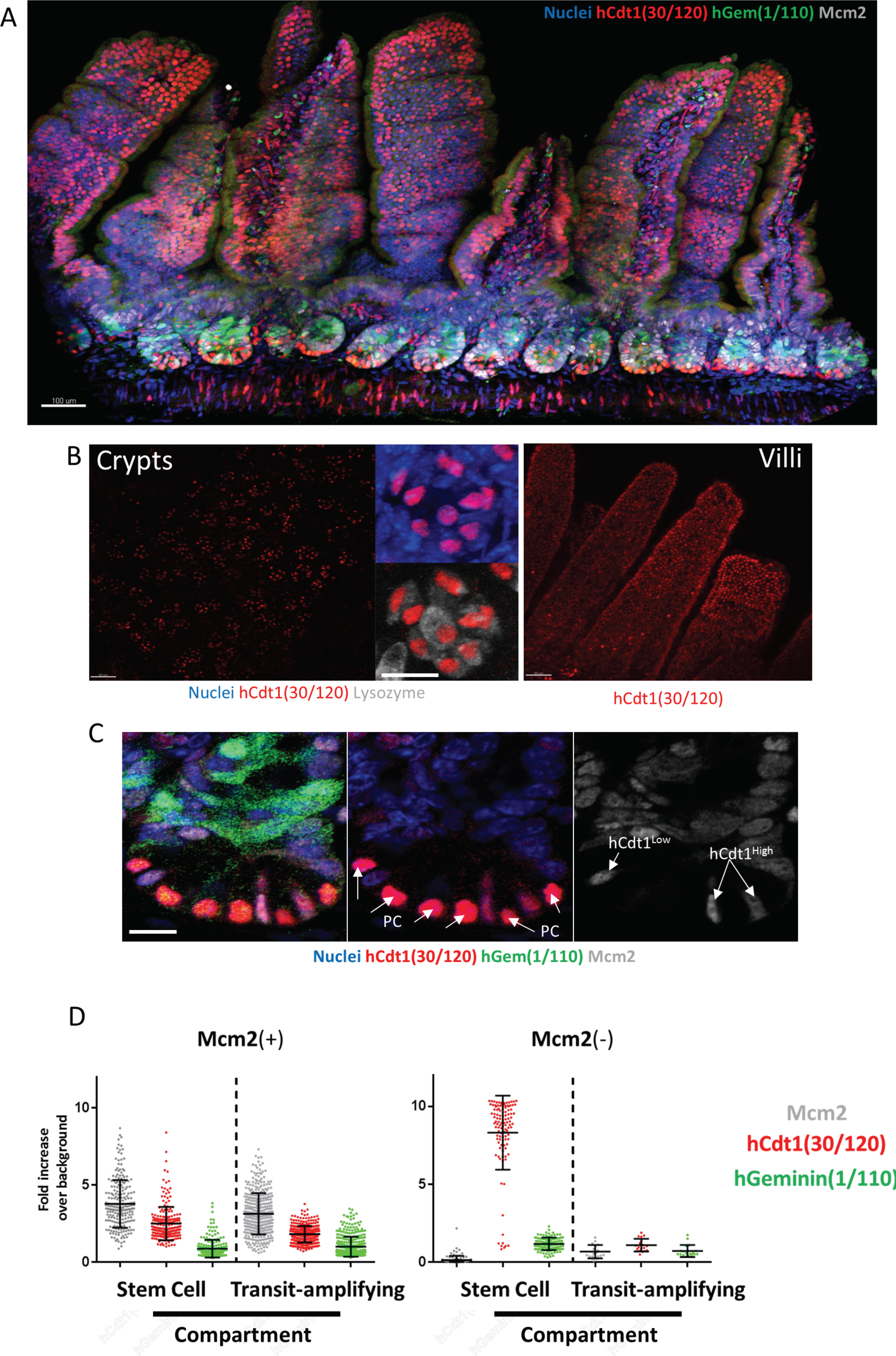
Delayed accumulation of Fucci2aR reporter hCdt1(30-120) in intestinal stem cells. (**A**) Representative image of a vibratome section of intestinal tissue derived from Fucci2aR mice, stained with Hoechst (Nuclei), an antibody against Mcm2 and showing expression of the G_1_ marker hCdt1(30-120) and S/G_2_/M marker hGeminin(1/110).
(**B**) Representative images hCdt1(30-120) expression in the crypt base (left panels) and villi (right panels). Most cells in the crypt base that highly express hCdt1(30-120) are UEA(+) Paneth cells.
(**C**) A representative image of a Fucci2aR intestinal crypt stained with Hoecsht and and antibody against Mcm2. All Mcm2(-) Paneth cells (PC) in the crypt base express high levels of hCdt1(30-120). Most Mcm2(+) stem cells in the crypt base express very low levels of hCdt1(30-120) (hCdt1^low^). Only few Mcm2(+) cells were found to express high levels of hCdt1(30-120) (hCdt1^High^). Many cells in the transit-amplifying compartment expressed hGeminin(1/110).
(**D**) Quantification of the expression of Mcm2, hCdt1(30/120) and hGeminin(1/110) in Mcm2(+) and Mcm2(-) cells in the stem cell and transit-amplifying compartment. Data is displayed as a ratio of signal to the background signal detected in negative cells (Stem cell compartment: Mcm2(+) N=220, Mcm2(-) N=112; Transit-amplifying compartment: Mcm2(+) N=383, Mcm2(-) N=18).

**S5 Figure.**
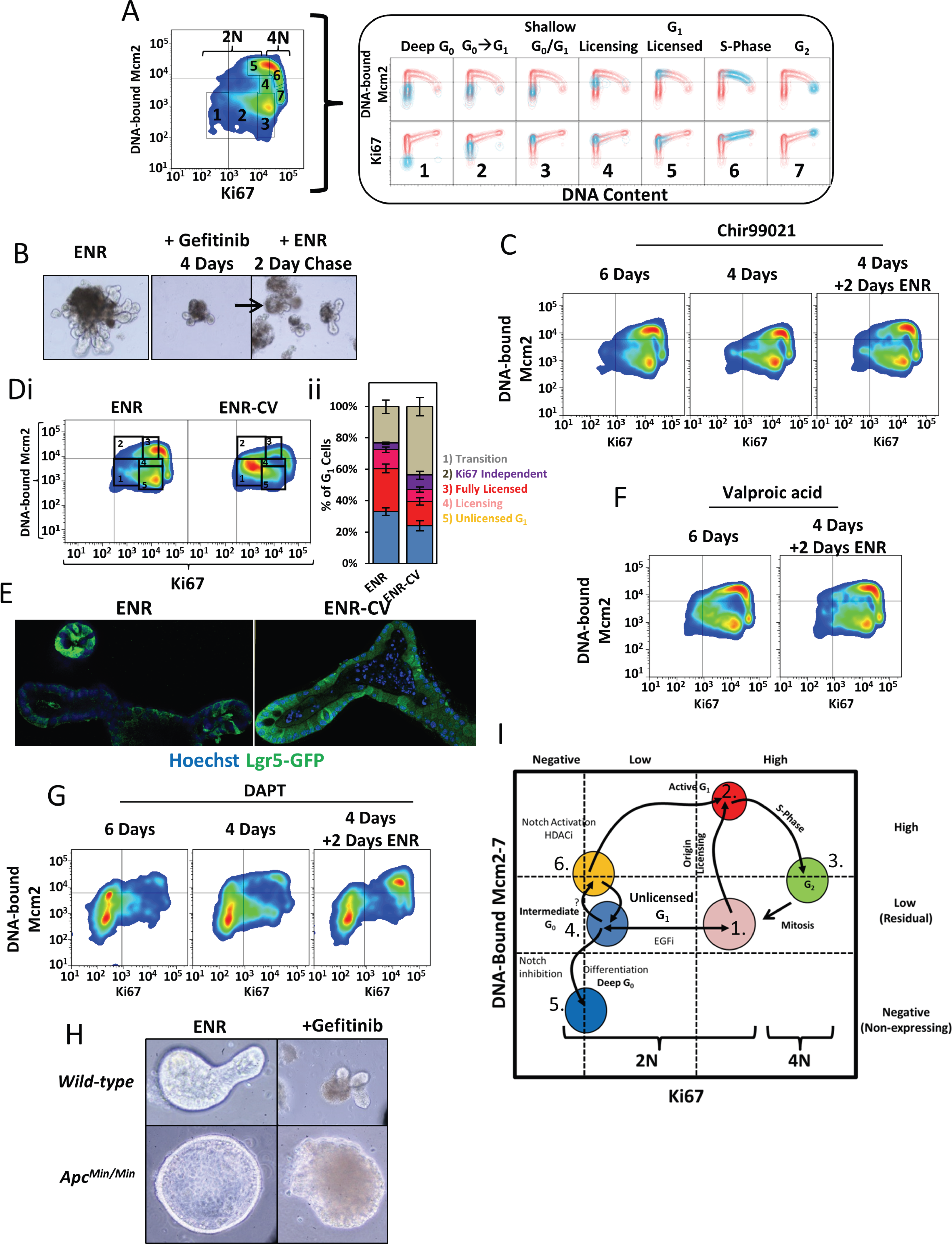
Manipulation of the stem cell niche can artificially induce Unlicensed-G_1_. (**A**) Representative flow cytometry profile of extracted epithelial cells isolated from organoids in ENR media. The displayed image is the same profile displayed in Figure 5C. The populations in boxed regions 1-7 are overlaid onto the individual Mcm2 and Ki67 cell-cycle profiles plotted against DNA-content (right hand box).
(**B**) Representative bright-field images of organoids in normal (ENR) media, treated with Gefitinib for 4 days, and after 2 days in fresh ENR media post Gefitinib treatment.
(**C**) Representative flow cytometry profiles from extracted cells isolated from organoids treated with 10μM Chir99021 for the indicated time intervals, and after 2 days in fresh ENR media post Chir99021 treatment.
(**D**) Representative flow cytometry profiles from extracted cells isolated from organoids treated with normal media (ENR) or in normal media containing Chir99021 and valproic acid (ENR-CV). Displayed is the comparison of DNA-bound Mcm2 vs Ki67 content (i). The gated populations: 1) Transition (Unlicensed-G_1_↔ intermediate G_0_), 2) Ki67 independent origin licensing pathway, 3) Fully licensed G_1_ cells, 4) Normal Licensing, 5) Unlicensed G_1_ have been quantified (N=3).
(**E**) Representative images of Lgr5-GFP organoids treated with ENR or ENR-CV. In ENR-CV treated organoids, the majority of cells express Lgr5.
(**F**) Representative flow cytometry profiles from extracted cells isolated from organoids treated with Valproic acid for indicated time intervals, and after 2 days in fresh ENR media post Valproic acid treatment.
(**G**) Representative flow cytometry profiles from extracted cells isolated from organoids treated with DAPT for indicated time intervals, and after 2 days in fresh ENR media post DAPT treatment.
(**H**) Representative images of *wild-type* and *Apc^Min/Min^* intestinal organoids after treatment with 5μM Gefitinib for 3 days.
(**I**) Model for the unique cell-cycle characteristics of organoid epithelial cells. Normal, highly proliferative cells, express Ki67 and Mcm2 protein that is not DNA-bound (**1**). During a normal cell-cycle, cells are activated from Unlicensed-G_1_, and rapidly license origins (**2**). Mcms are subsequently displaced during DNA replication (**3**) and remain unlicensed through G_2_ (**3**). Inhibiting EGFR causes highly proliferative cells (Ki67^hi^) to arrest in Unlicensed-G_1_ with maintained Mcm2 protein expression. Prolonged EGFRi treatment causes transition into an intermediate state of G_0_ accompanied by loss of Ki67 expression (Ki67^lo^), but maintenance of MCM2-7 protein expression (**4**). Induction of terminal differentiation by inhibition of Notch signalling is associated with a terminal loss of MCM2-7 proteins, and entry into deep-G_0_ (**5**). Notch activation (or HDAC inhibition (HDACi)) induces a unique subset of Ki67^lo^ Unlicensed-G_1_ cells to license origins independently of Ki67 **6**). We suggest that the unique cell population observed upon ENR-CV / ENR-V treatment may be a reserve subset of stem cells that express Lgr5 and start expressing MCM2-7 and enter Unlicensedμ from deep-G_0_. These cells have unique cell cycle characteristics, and can immediately license origins independently of Ki67 expression (6→2→3).

## S1 Movie. Cell-cycle clones

A 3D rotation of a representative intestinal crypt showing nuclei (Blue), DNA-bound Mcm2 (Red) and EdU (Green; 1 hour pulse). The crypt base is at the bottom of the image. An isosurface rendering of nuclei within the crypt has also been performed, and have been coloured to match specific cell-cycle stage: Unlicensed (Blue), G_1_ Licensed (Red), Early S-phase (Yellow), Late S/G_2_ (Green).

